# The genetic architecture of protein interaction affinity and specificity

**DOI:** 10.1101/2023.10.17.562688

**Authors:** Alexandra M. Bendel, André J. Faure, Dominique Klein, Kenji Shimada, Georg Kempf, Simone Cavadini, Ben Lehner, Guillaume Diss

## Abstract

Proteins function in crowded cellular environments in which they must bind to specific target proteins but also avoid binding to many other off-target proteins. In large protein families this task is particularly challenging because many off-target proteins have very similar structures. How this specificity of physical protein-protein interactions in cellular networks is encoded and evolves is not very well understood. Here we address the question of specificity-encoding by comprehensively quantifying the effects of all mutations in one protein, JUN, on its binding to all other members of a protein family, the 54 human basic leucine zipper transcription factors. Fitting a global thermodynamic model to the data reveals that most affinity changing mutations equally affect JUN’s propensity to bind to all its interaction partners. Mutations that alter the specificity of binding are much rarer. These specificity-altering mutations are, however, distributed throughout the JUN interaction interface. JUN’s interaction specificity is encoded by both positive determinants that promote on-target interactions and negative determinants that prevent off-target interactions. Indeed, about half of the specificity-defining residues in JUN have dual functions and both promote on-target binding and prevent off-target binding. Whereas nearly all mutations that alter specificity are pleiotropic and also alter the affinity of binding to all interaction partners, the converse is not true with mutations outside of the interface able to tune affinity without affecting specificity. Our results provide the first global view of how mutations in a protein affect binding to all its potential interaction partners and reveal the distributed encoding of specificity and affinity in an interaction interface. They also show how the modular architecture of coiled-coils provides an elegant solution to the challenge of optimising specificity and affinity in a large protein family.

## Introduction

Protein-protein interaction (PPI) networks form the backbone of a cell’s functional organization. The specificity of PPIs determines the structure of an interaction network and the way it coordinates cellular processes and thus plays an important role in establishing phenotypes. Specificity is determined by protein sequence and mutations that modify specificity rewire cellular networks, perturb the coordination of biological processes and modify organismal phenotypes, which can lead to disease, functional innovation and be subjected to natural selection. Understanding the physicochemical principles that govern how the sequence of a protein determines the specificity of its interactions is thus central to the question of how genotype determines phenotype.

Conserved families of protein interaction domains serve as models to study PPI specificity. One such family is the basic leucine zipper domains (bZIPs). The human genome encodes 54 bZIPs that form homo- and heterodimers that act as important transcription factors, including AP-1 (FOS/JUN), C/EBPs and MAFs. They are amongst the smallest known PPI domains and consist of an N-terminal basic DNA-binding domain of 27 amino acids and a C-terminal zipper domain of 35 amino acids that mediates homo- or heterodimerization via formation of coiled-coils^1–3^ (Fig. 1a). bZIP α-helices display a periodic pattern of seven amino acids per two turns. The zipper domains are therefore subdivided into five heptads within which the positions are labelled from *a* to *g*. Per heptad, side chains adopt the same position relative to the opposite bZIP in the dimer^4^. Amino acids at positions *a* and *d* are typically hydrophobic and form the hydrophobic core of the coiled coil interface by interacting with the facing amino acid at the same position in the partner, following a characteristic knobs-into-holes pattern that is essential for dimer stability and forms a zipper-like appearance from which its name is derived^1,5^. Positions *e* and *g* are often charged and can establish stabilizing salt bridges between the *g* residue of one helix and the subsequent *e* residue of the opposite helix^6^. Amino acids *b*, *c*, and *f* are solvent exposed in the coiled-coil and therefore often polar^4^.

**Fig. 1:**
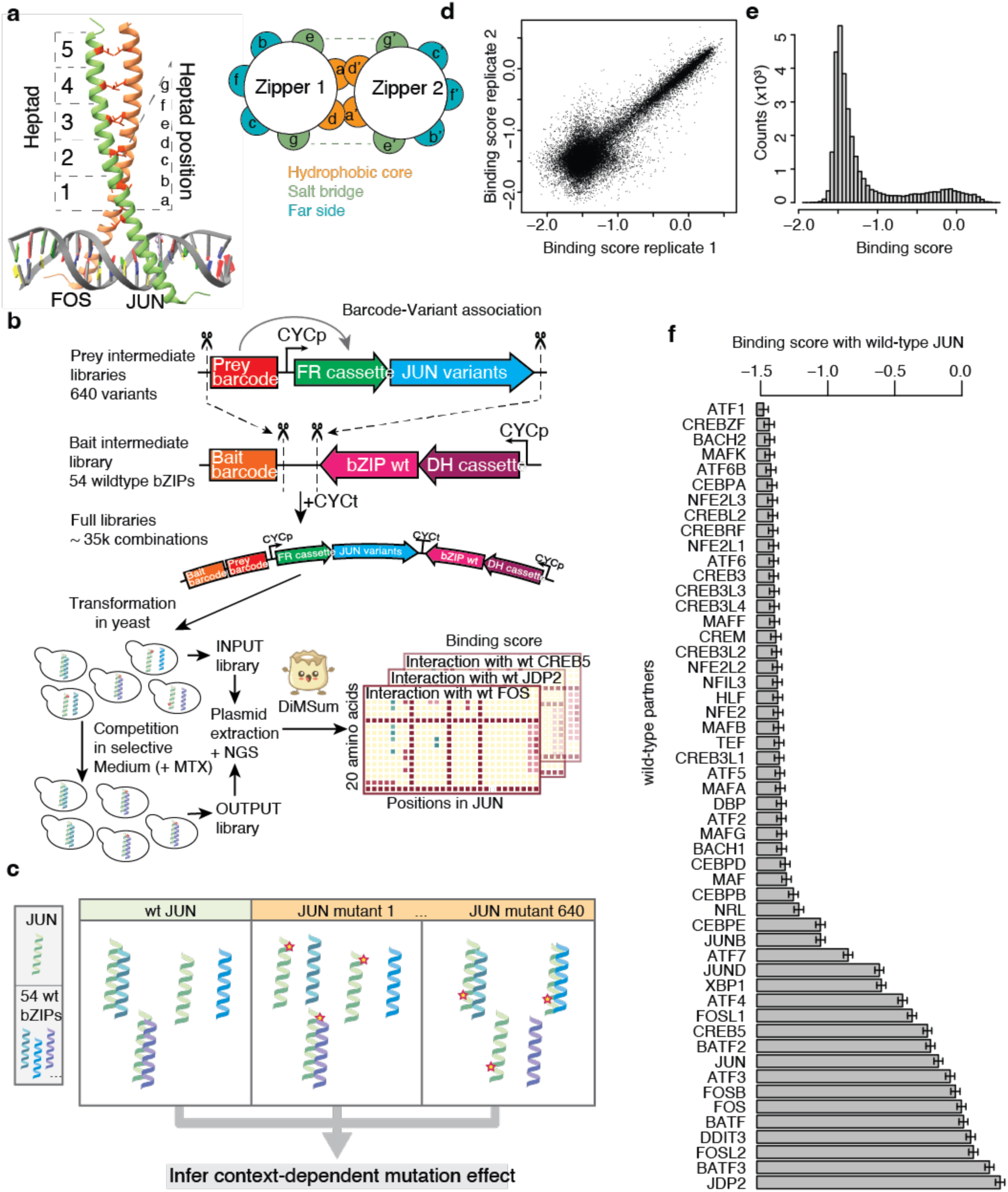
Experimental design. **a**, JUN-FOS dimer and heptad structure (pdb: 1fos). **b**, Cloning strategy and deepPCA. Two intermediate libraries were constructed. 640 variants of JUN were generated using overlap extension PCR and barcoded using random primer overhangs, 54 wild-type bZIPs were cloned and barcoded using random primer overhangs. The two intermediate libraries were cloned together to juxtapose the JUN variant barcodes with the wild-type bZIP barcodes. The full libraries were transformed into yeast cells, which were grown in selective medium containing Methotrexate. Enrichment of interacting pairs was quantified by sequencing barcode counts in INPUT and OUTPUT cultures by Next Generation Sequencing. Binding scores were calculated using DiMSum^23^ to assess the strength of interaction. **c**, Screening strategy. To identify context-dependent effects of mutations on the interaction between JUN and all 54 wildtype bZIPs, the effect of every possible single amino acid mutation was measured. This allowed the identification of determinants of specificity for individual interactions. **d**, Correlation between replicate 1 and 2 binding scores. **e**, Binding score distribution. **f**, Binding scores between wild-type JUN and all 52 wild-type partners. Error bars represent 95% confidence interval.

Our current understanding of bZIPs has mostly derived from the study of a small number of model leucine zippers, including AP-1 and the yeast Gcn4 homodimer^7–15^. Mapping bZIP interaction networks has revealed a sparse, highly modular network, with subfamilies of bZIPs displaying high specificities for one another and low levels of cross-talk^16^. However, this high level of specificity appears paradoxical with the simplicity of the coiled-coil interaction interface and the relative sequence similarity across bZIPs. This thus raises the questions of how a large set of high affinity interactions that use the same simple structural interface has evolved while maintaining high specificity and avoiding cross-talk or interference between the biological processes they regulate. Each bZIP binds only to a few other bZIPs. What are the determinants of this specificity, i.e. the residues that promote binding to one or a few other bZIPs and/or prevent binding to others? And how are these residues distributed within this simple interface?

Determining general, family-wide rules requires broader approaches where the effects of sequence variation are quantified not only on binding to one or a few known interaction partners but to all family members. We previously developed protein fragment complementation (PCA)-based deep mutational scanning assays to quantify the effects of hundreds of thousands of variants of two interacting proteins (Fig. 1b). These assays, which we collectively term deepPCA, are based on a split DHFR system whereby the proteins of interest are fused to complementary fragments of a murine DHFR variant (respectively called DH and FR) that confers resistance to methotrexate^17^. When the two proteins interact, the two fragments complement each other and the reconstituted DHFR activity sustains yeast growth in the presence of methotrexate, an inhibitor of the endogenous yeast DHFR. These assays are highly quantitative and have a large dynamic range^18–21^. In one deepPCA implementation, BindingPCA (bPCA), two proteins are expressed as DH and FR fusions from the same plasmid, enabling library-on-library screening in a single pool format (Fig. 1b, left). Yeast cells expressing pairs of proteins (wild-type or mutant) that form stronger interactions are enriched in the population while those forming weaker interactions are depleted. Deep sequencing then allows variants to be identified and the frequency of each variant or variant pair before and after growth in methotrexate to be quantified. The resulting enrichment scores are proportional to dimer concentration within the dynamic range of the assay and can be used to infer the underlying causal changes in binding free energy by fitting thermodynamic models to this data. bPCA thus provides a very high-throughput method to perform biophysical measurements^20–22^.

We previously employed bPCA to measure the effects of single and double mutants on the formation of the dimer between JUN and FOS bZIPs^20^. We showed that fitting a two-state thermodynamic model to the data improved our ability to predict how pairs of mutations combine and provided mechanistic insight into how mutations alter binding affinity. Our data showed that most combinations of genetic variants have additive effects on the free energy of binding that result in non-additive changes in dimer concentration because of the sigmoidal relationship between binding free energy and complex concentration. Double mutants not well predicted by the model identify energetically-coupled mutations, which we found to be enriched in directly physically contacting residues.

Here, we use bPCA to comprehensively quantify how single amino acid substitutions in JUN affect its binding to all 54 human bZIPs (Fig. 1c). The resulting data provide the first complete view of how changes in the sequence of a protein alter the specificity of its binding across an entire protein family.

## Results

### Deep mutational scanning of JUN’s complete bZIP interaction profile

To quantify the impact of substitutions on the binding specificity of JUN, we mutated each of 32 positions in the bZIP domain to every possible amino acid and quantified binding to all 54 human bZIPs (Table S1) using bPCA.

We improved bPCA by enabling the use of random DNA barcodes (Fig. 1b, left), which can be sequenced with shorter read lengths that are robust to sequencing errors, including those induced by template-switching (see methods). We constructed an intermediate library of JUN variants by overlap-extension PCR with NNS primers for each of the residues to be mutated and cloned them as FR fusions. Random barcodes were then added to each library, and the barcodes associated to each variant determined by deep sequencing (Table S2). We successfully obtained 615 out of the 640 total possible amino acid substitutions (96%; 32 positions x 20 substitutions, including stops), with a median of 38 different barcodes per substitution. The 54 human bZIPs were cloned as DH fusions, barcoded, verified by sequencing, and three barcodes per partner selected (Table S1). The final library was then assembled in a single pot reaction by combining the DH and FR intermediate libraries, juxtaposing the barcodes associated with the DH and FR fusions for efficient variant-bZIP identification by deep sequencing (Fig. 1b).

Binding of the JUN variants to all human bZIPs was assayed in six replicates and binding fitness scores and errors for each variant–partner pair calculated using DiMSum^23^ after collapsing read counts across barcodes associated to the same pair. Interaction scores were normalized to the wild-type binding of JUN and FOS (Table S3). After quality control, we were able to quantify the interaction of 26,648 out of the 34,614 expected pairs, including 579 unique variants and 52 out of the 54 wild-type bZIPs (excluding CEBPG and CREB1), with an average coverage of 443 reads per pair across input replicates. Binding scores are highly reproducible, with an average Pearson correlation between replicates of 0.93 (Fig. 1d).

The distribution of binding scores is bimodal (Fig. 1e), with a smaller number of variant-partner pairs in the high binding score mode, reflecting the relatively small fraction of high affinity interaction partners that wild-type JUN has (Fig. 1f). As previously reported, wild-type JUN forms strong interactions with many members of the AP-1 family, including BATF1-3, the FOS subfamily, the JUN-subfamily, JDP2, ATF3, 4 and 7. We also detected previously reported interactions with CREB5 and DDIT3^16,24^ and a previously unreported interaction with XBP1, a regulator of the unfolded protein response^25^.

### Most mutations similarly affect binding to all interaction partners

In order to compare the effect of the same mutation with different partners, we normalized each measurement by the interaction between the corresponding human bZIP and the JUN wild-type allele. Fig. 2a shows the relative effects of all JUN mutants on the interaction with all bZIPs. The partners are ordered according to their binding score with wild-type JUN, revealing a general pattern of mutational effects that is highly similar across all JUN interaction partners. This pattern identifies positions that are highly sensitive to mutation, such as the *a* and *d* positions of all heptads, consistent with them forming the hydrophobic core of the interaction interface^3^. These mutations affect binding to all of JUN’s partners in a similar way, i.e. they affect the intrinsic capacity of JUN to dimerize with other bZIPs. For example, replacing one of the core leucines that define bZIPs by a charged amino acid disrupts JUN’s ability to form bZIP dimers regardless of the partner (Fig. 2a).

**Fig. 2:**
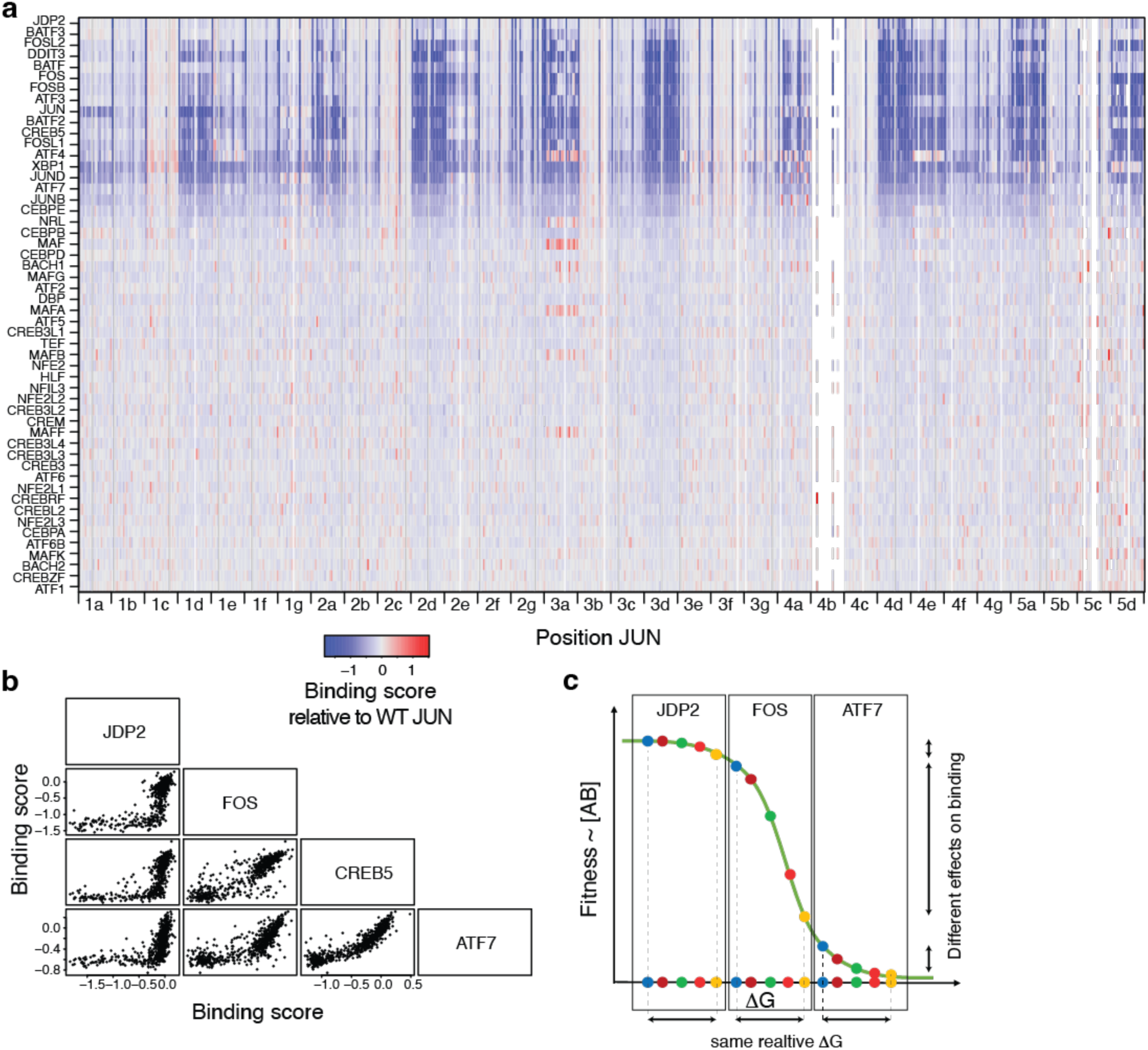
Mutational effects are mostly non-specific. **a**, Heatmap of binding scores relative to the corresponding interaction between the wild-type partner and wild-type JUN. Rows are ordered by decreasing strength of the wild-type interaction. **b**, Binding score profile of four selected partners, showing a non-linear trend between mutations that correspond to general effects of mutation. **c**, The non-linearity trend between mutational profiles is easily explained by the non-linearity between ΔG and binding scores if mutations have a general effect at the energetic level that is independent of the partner.

Although most of the mutational effects appear qualitatively as having a general effect on JUN’s ability to dimerize, some mutations appear to affect only the interaction with some partners. These represent cases of specific effects. For instance, several substitutions at position *3a* (position *a* in heptad 3) increase the interaction of JUN with members of the MAF subfamily (Fig. 2a).

Plotting the mutational effects profiles across partners also reveals a strong trend affecting all partners, showing that most mutations have general effects on JUN’s ability to form dimers with all its partners (Fig. 2b). The non-linearity of these relationships is reminiscent of global effects due to the thermodynamics of binding to partners with different wild-type affinities (Fig. 2c). Some mutations, however, change binding to particular interaction partners such that their binding score is located away from the general trend. These residuals represent mutational effects that are specific to these partners. To formally quantify these changes in binding specificity we therefore used modelling to decompose mutational effects into their general and specific components while accounting for the non-linearity imposed by the thermodynamics of binding.

### A global two-state additive thermodynamic model accurately predicts changes in binding to 52 bZIPs

Using MoCHI^21,22^, we fitted a simple two-state thermodynamic model to the data where proteins cooperatively fold upon binding and so can exist in an unbound, unfolded state or in a folded and bound state (Fig. 3a). MoCHI infers a single additive binding Gibbs free energy change (ΔΔG) for each of the 640 JUN mutations and 51 additive binding free energies for each partner bZIP (relative to FOS as an arbitrary reference, see methods). A single ΔΔG, which we refer to as the change in global binding energy, is fitted for each variant or partner across the entire dataset. MoCHI assumes additivity for all changes in free energy (ΔΔG) and a linear relationship between the fraction of bound molecules and the binding scores quantified by bPCA^19–21^. Therefore, the fitted variant ΔΔGs capture the general effects of mutations across all partners. The residuals between the experimentally measured effects of mutations and the model predicted values then quantify the specific effects of mutations with each binding partner.

**Fig. 3:**
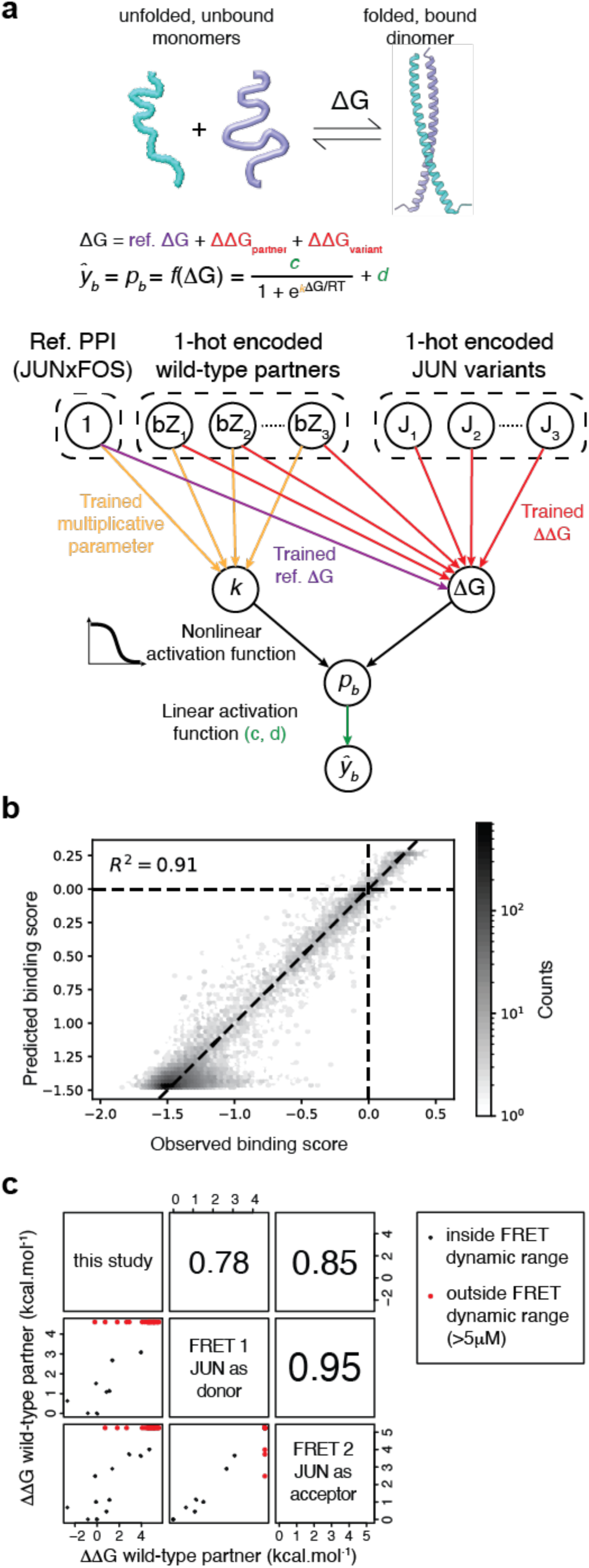
A simple thermodynamic model predicts well mutational effects. **a**, MoCHI^21,22^ fits a simple two-state thermodynamic model between unbound, unfolded and bound, folded states. The total DG corresponds to the sum of the 1′G of the reference protein interaction, arbitrarily chosen as wild-type JUNxFOS, the ΔΔG of the mutation relative to wild-type JUN, and the ΔΔG of swapping FOS for another bZIP. MoCHI uses one-hot encoded sequence representing which substitution at which position in JUN is present and a dummy sequence representing which bZIP partner is present. It then fits the reference 1′G, the mutant ΔΔG and the partner ΔΔG and sum them up, and in parallel a multiplicative parameter for each partner that corrects a small bias in the distribution of residuals due to different curvature of the sigmoid for different partners and that relates to a Hill coefficient (see methods). **b**, Plot of observed binding scores versus the ones predicted by the thermodynamic model. **c**, ΔΔG inferred by MoCHI correlate well with those determined by *in vitro* FRET^24^ whether JUN was used as FRET donor or acceptor (n=34).

The additive MoCHI energy model predicts binding scores with high accuracy (R^2^ = 0.91, Fig. 3b), and the partner ΔΔGs correlate well with independently-measured *in vitro* binding energies^24^ (Pearson’s R = 0.78 and 0.85 when JUN is labelled either as donor or acceptor in fluorescence resonance energy transfer (FRET) experiments, respectively, n=34; Fig. 3c), confirming the validity of our approach. Some well-known JUN binding partners appear below the detection limit of the *in vitro* FRET assay, for instance JUNB or the JUN homodimer, suggesting these correlations may underestimate the agreement of MoCHI’s ΔΔGs with the ground truth. The high correlation between the measured binding scores and those predicted using an additive energy model in which the effects of mutations are the same for all interaction partners shows that the vast majority of measured changes in binding are independent of the interaction partner’s identity and that context-dependent, specific effects are rare.

### Global binding energy changes are determined in part by helical propensity and interface hydrophobicity

To better understand the underlying mechanistic causes of changes in global binding energy, we used linear regression to predict the fitted ΔΔGs from a compilation of ∼500 physicochemical properties of the 20 natural amino acids^26^, which heptad is mutated and the position within the heptad. The five top features together explain 68% of the variance in non-specific effects (Fig. 4a, b), with the top feature related to alpha helical stability^27^, the two features encoding heptad number and position within the heptad, and the last two features related to side chain hydrophobicity^28^ and standard entropy^29^ (related to number of possible conformations), respectively. Thus, mutations decreasing the stability of JUN’s alpha helix and/or its hydrophobicity tend to reduce its intrinsic ability to interact with other bZIPs, regardless of the partner^2,30,31^.

**Fig. 4:**
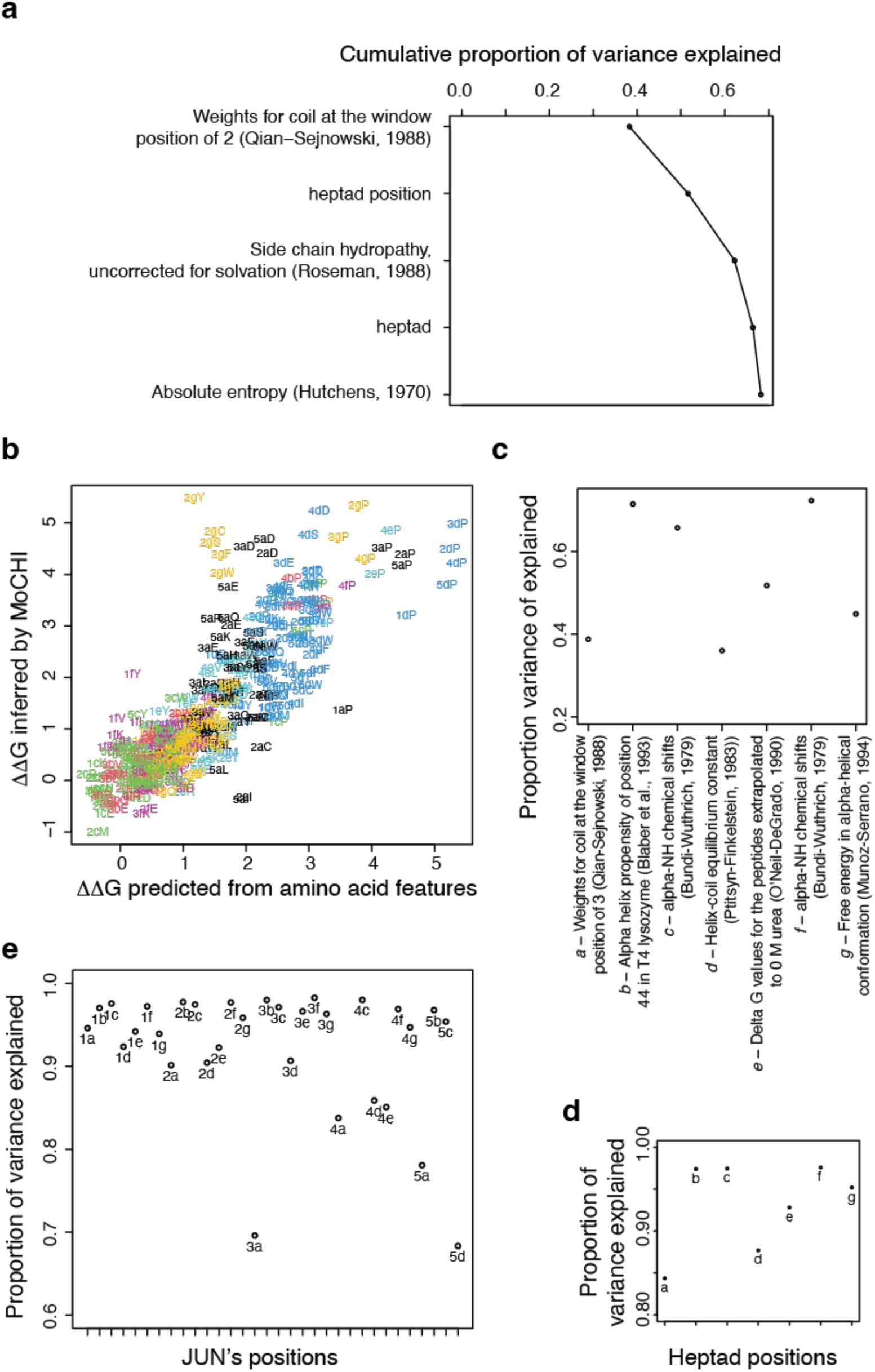
The general component of mutational effect corresponds to JUN’s propensity to form a bZIP interface. **a**, Cumulative proportion of variance in mutant ΔΔG explained by the five best features, including three physicochemical properties of amino acids related to alpha-helical stability, hydrophobicity and conformation. **b**, Mutant ΔΔG predicted by amino acid features. X-axis corresponds to the ΔΔG predicted by the five top features represented in panel **a**, and y-axis corresponds to the ΔΔG fitted by MoCHI. **c**, Proportion of variance in mutant ΔΔG explained by the top feature at each of the seven heptad positions. **d,e** Proportion of variance in binding scores explained by global binding energy at each of the seven heptad positions (**d**) and each of the 32 mutated positions (**e**).

Because of the importance of position in the model, we next looked at the most predictive feature at each of the seven heptad positions independently (Fig. 4c). Far-side positions *b*, *c* and *f* were best predicted (best feature explaining 71, 66 and 72% of the variance, respectively)^32,33^, followed by salt-bridge positions *e* and *g* (52 and 45% of the variance, respectively)^34,35^, and hydrophobic core positions last (39 and 36% of the variance explained for *a* and *d*, respectively)^27,36^.

A substantial part of the changes in global binding energy inferred by the model therefore reflects the general ‘zipper propensity’ of a sequence i.e. the baseline ability to form a coiled coil and dimerize with other bZIPs.

### Global binding energy changes explain most mutational effects outside of the interface

We compared the variance in binding explained at each of the seven heptad positions by changes in the global binding energy. Positions on the far side of the helix (*b*, *c* and *f*) have the highest proportion of variance explained by global binding energy while hydrophobic core positions (*a* and *d*) have the lowest (Fig. 4d, e). This is in agreement with core positions making direct contacts with binding partners.

### Changes in binding specificity

Plotting the binding scores against the global binding energies for each bZIP partner (Fig. 5a, Fig. S1) reveals that global binding energies predict the binding fitness estimates very well for all bZIPs except ATF4 (which is removed from subsequent analyses, see methods). However, for each bZIP, some changes in binding are not well predicted by change in global binding energy, with large residuals from the predicted binding. These residuals quantify changes in binding specificity, i.e. mutational effects that are specific to one or a subset of interaction partners. Inspection of these plots suggests these specificity-altering mutations are enriched in specific positions and for particular types of substitutions for each interaction partner (Fig. 5a).

**Fig. 5:**
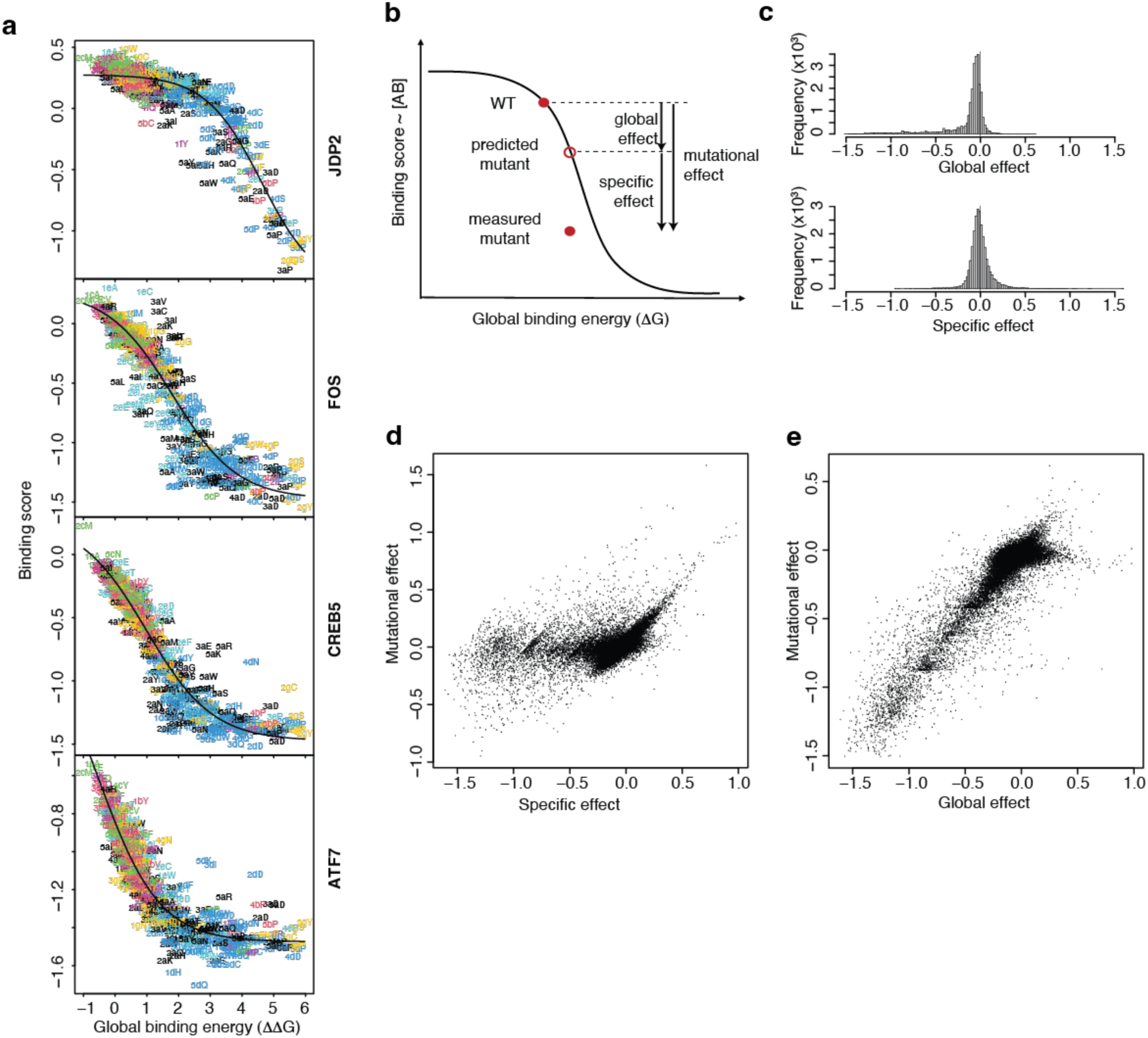
Mutational effects can be decomposed in their general and specific component. **a,b**, Specifcity plots representing the binding scores as a function of the global binding energy (ΔΔG) inferred by MoCHI. The sigmoid curve represents binding scores predicted by MoCHI, and the vertical residual from the curve thus represent the specific effect of this mutation on the interaction with this partner. The three characters in the mutant name corresponds to heptad, position within the heptad and amino acid substitution, respectively. Mutant names are colored according to their heptad position, which further highlights that several mutations at the same few positions have specific effects. Examples with the four partners presented in Fig 2B (**a**) and general principle of the decomposition of specific and global effects (i.e. effects due to change in global energy) (**b**). **c**, distribution of mutational effect du to changes in global binding energy (top) and specificity (bottom). **d,e**, Plot of mutational effects as a function of their specificity (**d**) and global binding energy (**e**) components. Each point represents a different variant-partner pair.

The energy model therefore allows us to quantitatively decompose the measured change in binding caused by each mutation for each interaction partner into global and specific components. We define the global component of mutational effects as the difference between the mutant binding score predicted by the thermodynamic model and the wild-type binding score, and the specific component as the difference between the measured mutant binding score and the predicted one (Fig. 5b). The distributions of these effects are rather different, with global effects having a long negative tail and specificity changes having a more pronounced positive tail (Fig. 5c). Out of all variant-partner pairs tested, 8887 interactions are altered by a change in global binding energy (global effect ≠ 0; FDR < 5%), while only 207 are altered by a change in specificity (specific effect ≠ 0; FDR < 5%). In summary, mutations are much more likely to affect JUN’s affinity globally than specifically.

Plotting changes in binding as a combination of changes in specificity and global energy changes reveals that decreased binding is mostly caused by changes in global energy whereas increased binding is mostly due to changes in specificity (Fig. 5d,e).

### Negative, positive and dual determinants of interaction specificity

Clustering the specificity profile of each bZIP reveals that members of the same subfamily tend to cluster together, especially for those that interact strongly with JUN, and identifies the wild-type positions important to establish specific interactions with a given subfamily (Fig. 6a). We next quantitatively characterize these determinants of specificity and how they are distributed throughout the bZIP interface.

**Fig. 6:**
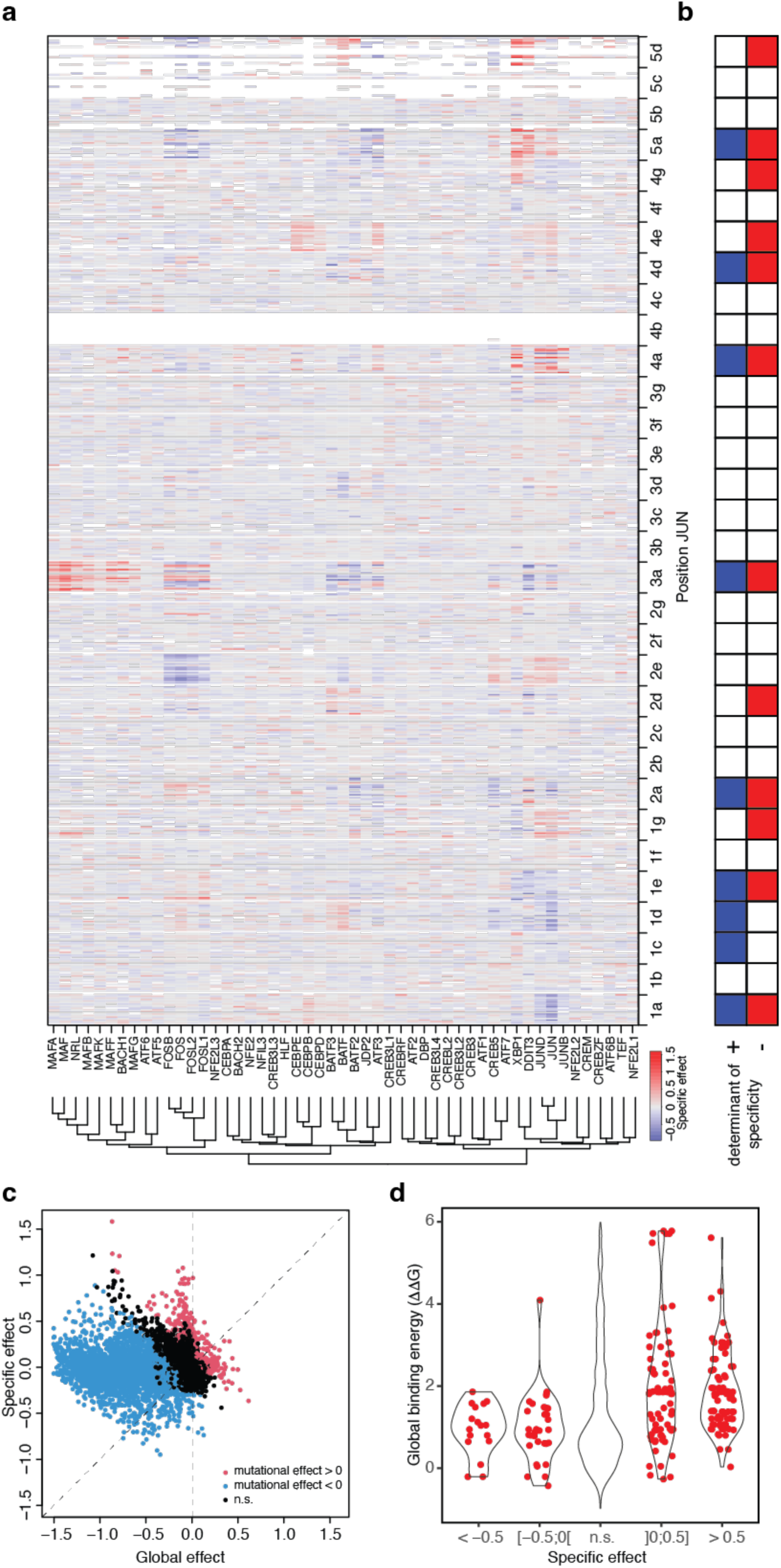
JUN’s determinants of specificity. **a**, Heatmap of specific mutational effects revealing clusters of mutations at a few positions that affect some partners specifically. Columns are clusters according to each partners specificity profile. **b**, Summary of positions involved in determining JUN’s specificity. Positions were labelled as a positive or negative determinant of specificity if it had at least one significant specific mutational effect with one partner (FDR < 5%, specific effect < -0.5 or > 0.5, respectively). **c**, Relationship between the specific and global binding energy components of mutational effects. Each point represents a different variant-partner pair and is colored according to its mutational effect relative to the binding score of the corresponding wild-type interaction (FDR < 5%). **d**, Changes in global binding energy for different bins of specific effects, showing the pleiotropic nature of specificity-changing mutations (FDR < 5%).

Binding specificity can potentially be determined via three different strategies. Wild-type residues can be positive determinants of specificity that increase affinity to one or a subset of bZIPs, negative determinants that decrease affinity to one or a subset of bZIPs, or dual determinants that both promote on-target binding and prevent specific off-target binding^37–39^. Our comprehensive mutagenesis and mutational effect decomposition allows us to quantify the relative prevalence of each strategy for the first time in a large protein family.

When mutated, positive determinants of specificity result in reduced binding to only some interaction partners. Conversely, mutation of negative determinants leads to increases in binding to only some interaction partners. Mutating dual determinants of specificity leads to positive effects with some partners and negative effects with others.

Out of the 97 variant-bZIP combinations with strong changes in specificity (absolute specific effect > 0.5, FDR < 5%), 80 are positive and 17 negative (82.5% and 17.5%, respectively). These represent a total of 56 unique mutations out of the 579 tested (9.7%), with 41 having only positive effects, 11 only negative effects and 4 both positive and negative effects. 16 partners are affected by positive effects, with an average of 4.6 positive mutations per partner. Only five partners are affected by negative effects, with four of them affected by a single negative mutation and eight mutations affecting JUN binding with itself. Dual mutations affect nine partners, with an average of 1.33 dual mutations per partner.

However, some mutations with strictly negative or positive effects are found at the same position, such that the wild-type residue can be considered a dual determinant of specificity, with different mutations having different directions of effect. Negative determinants of specificity thus correspond to positions where mutations only have positive effects and positive determinants of specificity correspond to positions where mutations only have negative effects. Out of the 32 tested positions in the zipper domain, seven are dual determinants of specificity, five are negative determinants and two are positive determinants (Fig. 6b).

The interaction specificity of JUN is thus defined by a mixture of negative and positive specificity determinants, with half of the specificity determining positions acting both negatively and positively to encode specificity.

### Positive, negative and dual determinants of specificity are dispersed throughout the interaction interface

We next examined where in the JUN sequence the determinants of specificity are located. Across the five heptads, residues in five out of seven positions (*a*, *d*, *c*, *e*, *g*) contribute to specificity. Position *a* in all five heptads is a dual determinant of specificity, as is position 1e and 4d (Fig. 6b). Negative determinants of specificity are found at position 1g, 2d, 4e, 4g and 5d, while positive determinants are found at position 1c and 1d. The strictly positive determinants of specificity are thus more biased towards the N-terminal region of the leucine zipper while the strictly negative determinants are more biased towards the C-terminal region.

Interestingly, three out of the four salt-bridge positions that contribute to specificity act negatively, with the other being a dual determinant. JUN therefore employs electrostatic interactions mostly in a repulsive rather than attractive manner to determine specificity.

### Mutations affecting specificity are nearly always pleiotropic and also affect the global binding energy

We next examined whether changes in specificity also affect the global binding energy. Mutations that alter specificity might have no effect on the global binding energy (if specificity and global binding energy are orthogonally encoded) or mutations causing specificity changes may have pleiotropic effects and synergistically or antagonistically alter affinity via changes in both specificity and global binding energy.

We find that in JUN 50 out of 56 (89.3%) unique mutations that change specificity to at least one partner also change the global binding energy (Fig. 6c, d). In nine cases, changes in specificity and global binding energy have synergistic effects on affinity (all negative) and in 37 cases changes in specificity and global binding energy have antagonistic effects on affinity. Out of these, a single mutation has a negative effect on specificity and positive effects on global affinity, while 36 have positive effects on specificity and negative effects on global affinity. The last four mutations have both negative and positive effects on specificity, depending on the interaction partner and in all cases affecting negatively the gobal affinity (Fig. 6c, d).

We conclude that nearly all mutations affecting specificity are pleiotropic and also alter binding affinity to all other interaction partners. However, the reverse is not true and the vast majority of mutations that alter the global binding energy do so without changing the binding specificity.

### Hydrophobic core positions in the first and last heptad are important determinants of specificity

Core positions *a* and *d* are important for establishing specificity, with strong specific effects for all *a* positions across the five heptads and for *d* positions in the extremity heptads 1 and 5 (Fig. 6a). Mutations at position 1d have a relatively small absolute effect on interaction with wild-type partners compared to other *d* positions (Fig. 2a). For most of JUN’s wild-type partners, two of the surrounding positions in the hydrophobic core, 1a and 2a, are bulky or hydrophilic (Fig. 7a, left), likely weakening the contribution of the 1d leucine and reducing the overall importance of the first heptad to binding. However, JUN subfamily members have hydrophobic residues at 1a and 2a (Fig. 7a, right), making heptad 1 important for binding such that mutations in JUN positions 1a and 1d have specific effects.

**Fig. 7:**
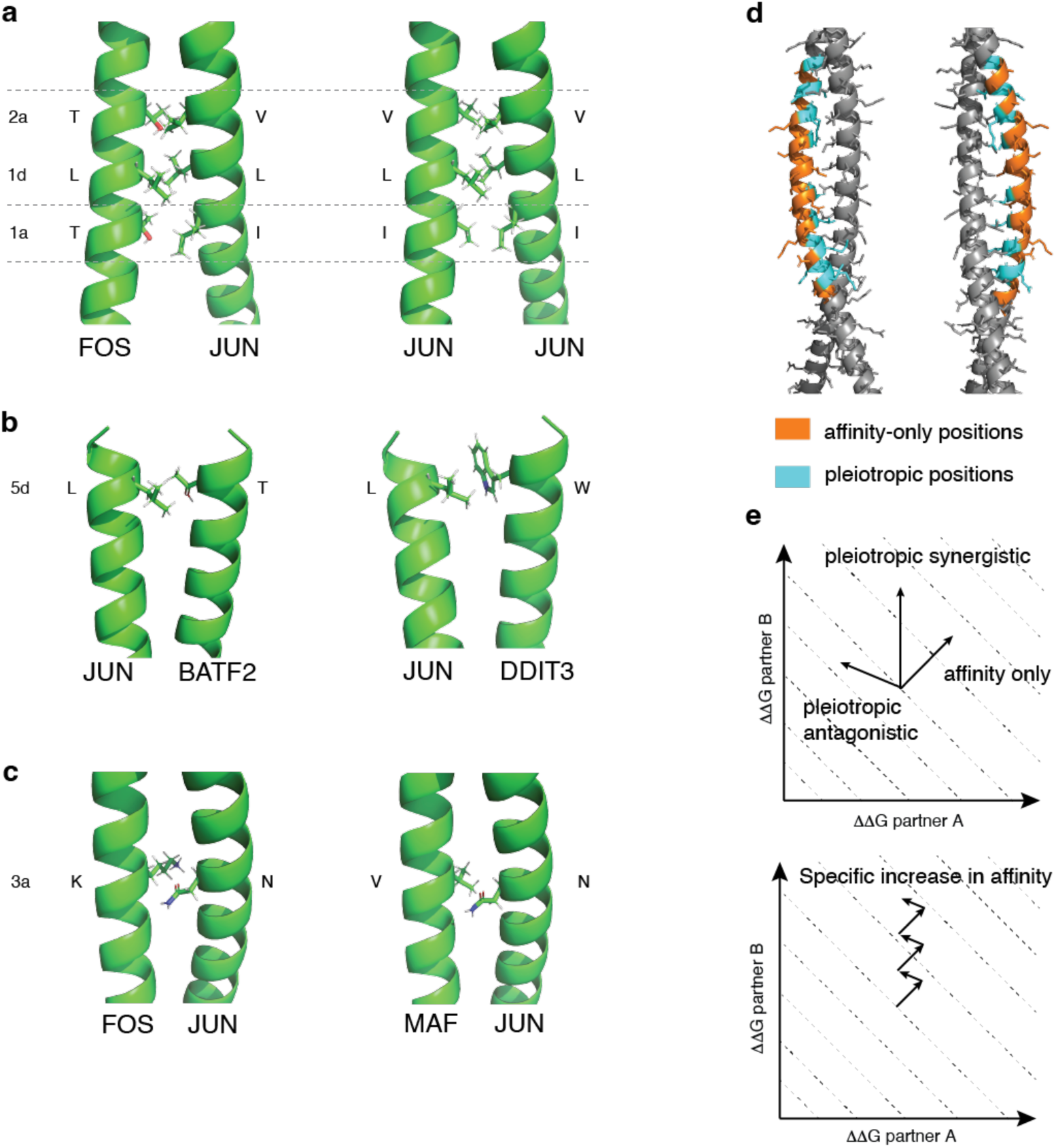
Structural insights into the mechanisms of the determinants of specificity. **a**, **b**, **c**, structural insights into a few examples of determinants of specificity. All structures were predicted using AlphaFold2^46^ since crystal structures are not available for many of the dimers. **d**, Distribution of affinity-only and pleiotropic positions within the bZIP interface. Crystal structure of the JUN-FOS dimer^49^. **e**, zig-zag model of bZIP evolution. Because of the pleiotropic antagonistic nature of nearly all mutation changing specificity, evolution of specificity requires compensatory mutation at the far side increasing affinity globally by changing global binding energy only.

Mutations at positions 5d generally weaken the interaction with all partners, as expected for a core leucine, but they do so less than expected for the BATF subfamily as well as for DDIT3 and XBP1, even in some cases making the interaction stronger than with wild-type JUN (Fig. 6a). BATF and BATF2 have bulky aromatic residues at the facing 5d position (Fig. 7b, left), which could explain why mutating JUN towards alanine, valine and isoleucine, all less bulky than leucine (isoleucine has the same number of carbon atoms, but one is closer to the backbone than in leucine), are amongst the most positive residuals for these partners. For interaction with DDIT3 and XBP1, nearly all 5d substitutions have positive specific effects, with a large fraction increasing the interaction with XBP1 compared to wild-type JUN (Fig. 6a). The presence of a threonine at position 5d in DDIT3 (Fig. 7b, right) and at position 5a in XBP1 suggests a weakened hydrophobic core in this last heptad, such that mutating the leucine has a reduced impact on the contribution of this heptad hydrophobic core to binding compared to other partners that have a 5d leucine.

### Asparagine at position 3a is a trade-off between preventing binding to the MAF subfamily and promoting binding to the FOS and BATF sub-families

Another interesting position is 3a. The wild-type asparagine at position 3a is involved in the specificity of the interaction with the FOS-subfamily since some mutations have negative specific effects. However, asparagine is not the optimal residue for FOS subfamily binding with mutations to valine and cysteine increasing binding. However, although valine is the optimal amino acid at this site for FOS subfamily-binding it also increases binding with the MAF subfamily (Fig. 6a). MAF bZIPs have a complementary valine at the facing 3a position. Asparagine at position 3a may thus reflect an evolutionary trade-off between promoting the interaction with the FOS subfamily and avoiding strong interactions with the MAF subfamily.

## Discussion

We have presented here the first comprehensive quantification of the effects of mutations in a protein on its binding to possible interaction partners from a protein family. Our data quantifies the binding of JUN to 52 of the 54 members of the human bZIP family of proteins: a total of 26,648 measurements. Fitting a global thermodynamic model to this data allowed us to deconvolve the effects of mutations on binding specificity from changes in the propensity of JUN to bind to all its interaction partners (Fig. 2a, b & Fig. 5a, b).

Our data show that mutations overwhelming affect JUN’s binding to interaction partners non-specifically (Fig. 5c, e). Mutations in all 32 tested positions can alter this global binding energy, which, at least in part, relates to both the helical propensity of the JUN sequence and the hydrophobicity of the interface (Fig. 4).

Mutations that alter the specificity of binding are rarer (Fig. 5c), but they are also distributed throughout the protein interface (Fig. 6a, b & Fig. 7e). Indeed, mutations in nearly half of amino acid positions alter JUN’s binding specificity. The core interface residues *a* and *d* are particularly important determinants of specificity. The binding specificity of JUN is defined by a mixture of both positive determinants that increase the affinity for target proteins and negative determinants that reduce binding to off-target proteins. Whilst the positive determinants are enriched in the C-terminus, the negative determinants are more abundant in the N-terminus of the zipper. The negative specificity determinants are both larger in number and individually of larger effect (Fig 5c). Preventing binding, for instance through repulsive electrostatic interactions, is therefore a more important strategy for defining specificity than promoting binding. However, half of the specificity-defining residues serve as both positive and negative determinants of JUN’s interaction profile. Specificity is thus encoded in a delocalized manner and by a mixture of positive and negative interactions, with many residues acting as dual specificity determinants.

Specificity and affinity in JUN are, however, not encoded by orthogonal genetic variables. Our data show that it is difficult to change the specificity of JUN without also altering its affinity for all interaction partners (Fig. 6c, d). Put another way, specificity-altering mutations are nearly always pleiotropic and also change the affinity of the protein for all of its interaction partners. During evolution, therefore, changes in specificity will rarely occur without changes in affinity to all interaction partners. Specificity changes will therefore normally have to be accompanied by compensatory mutations that counteract the change in affinity without altering specificity.

However, although changes in specificity nearly always also involve changes in affinity to all binding partners, the reverse is not true and there are many mutations in many positions that can alter the affinity of JUN for all interaction partners without altering its specificity. The modular design of coiled coils thus provides an elegant solution to the problem of independently tuning specificity and affinity in a large protein family, allowing both properties to be optimized. Whereas mutations in the interface affect both specificity and global affinity, mutations outside of the interface alter the general zipper propensity and so affinity without affecting specificity (Fig. 7d). Specificity can therefore be optimised during evolution using combinations of pleiotropic specificity and affinity altering mutations in the interface coupled to compensatory affinity tuning mutations elsewhere in the zipper (Fig. 7e). We refer to this model as the zig-zag model of specificity and affinity optimisation.

Our results concern a particular protein – JUN – from a particular family of proteins – basic leucine zippers – and a particular class of protein interactions – coiled coil interactions. In future work it will be important to use similar experimental designs to better understand the encoding of affinity and specificity in other protein families. Pioneering studies have already begun to address these questions for other structural classes of protein interactions^40–42^ and we believe that a concerted effort to mutagenize the diversity of domain-peptide and domain-domain interactions that constitute the interactomes of every species will allow us to better understand, predict and engineer affinity and specificity from sequence.

## Methods

All plasmids and strains are available upon request.

### Yeast Strain

All experiments were performed in BY4742 (MATα *his3*Δ1 *leu2*Δ0 *lys2*Δ0 *ura3*Δ0).

### Media and buffer recipes

- LB: 25 g/L Luria-Broth-Base (Invitrogen™). Autoclaved 20 min at 121 °C.
- LB-agar with 2X Ampicillin (100 μg/mL): 25 g/L Luria-Broth-Base (Invitrogen™), 7.5 g/L Agar, 1.2 g/L MgSO_4_ · H_2_O. Autoclaved 20 min at 121 °C. Cool-down to 45 °C. Addition of 100 mg/L Ampicillin.
- YPAD: 20 g/L Bacto-Peptone, 20 g/L Dextrose, 10 g/L Yeast extract, 25 mg/L Adenine. Filter-sterilized (Millipore Express ®PLUS 0.22 μm PES, Merck, Darmstadt, Germany).
- SC-ura: 6.7 g/L Yeast nitrogen base without amino acids, 20 g/L glucose, 0.77 g/L complete supplement mixture drop-out without uracil. Filter-sterilized (Millipore Express ®PLUS 0.22 μm PES, Merck, Darmstadt, Germany).
- SC-ura/ade/met: 6.7 g/L Yeast nitrogen base without amino acids and folic acid, 20 g/L glucose, 0.74 g/L complete supplement mixture drop-out without uracil, adenine and methionine. Filter-sterilized (Millipore Express ®PLUS 0.22 μm PES, Merck, Darmstadt, Germany).
- SORB: 1 M sorbitol, 100 mM LiOAc, 10 mM Tris-HCl pH 8.0, 1 mM EDTA pH 8.0. Filter-sterilized (Millipore Express ®PLUS 0.22 μm PES, Merck, Darmstadt, Germany).
- Plate mixture: 40 % PEG3350, 100 mM LiOAc, 10 mM Tris-HCL pH 8.0, 1 mM EDTA pH 8.0. Filter-sterilized (Millipore Express ®PLUS 0.22 μm PES, Merck, Darmstadt, Germany).
- Recovery medium: YPAD + 0.5 M sorbitol. Filter-sterilized (Millipore Express ®PLUS 0.22 μm PES, Merck, Darmstadt, Germany).
- Competition medium: SC-ura/ade/met + 200 μg/mL methotrexate (BioShop Canada Inc., Canada), 2 % DMSO.
- DTT buffer: 0.1 M EDTA-KOH pH7.5, 10 mM DTT
- Zymolyase buffer: 20 mM K-phoshpate pH 7.2, 1.2 M sorbitol, 0.4 mg/mL Zymolyase 20T (amsbio, USbiological), 100 μg/mL RNAse A

### Plasmid backbone construction

Two intermediate plasmids were constructed, one carrying the DHFR-bait cassette, one carrying the DHFR-prey cassette. For the bait intermediate plasmid, a gene fragment carrying the CYC promoter, a 3 x Myc-tag fused to a DHFR N-terminal half (from here on referred to as DH-tag), a Linker sequence (Linker L3, GSAGASAGGSGSAGSASGGS), a 34 bp random spacer sequence, a 20 bp placeholder barcode sequence (GGCACTGTAGTCGATAGCCT; bait barcode) and an SP1 Illumina primer binding site (Illumina, San Diego, CA) was cloned in pRS416^43^ digested with KpnI-HF (New England Biolabs, Ipswitch, MA, USA) and SacI-HF (New England Biolabs, Ipswitch, MA) using Gibson assembly. The Gibson assembly was performed using 34 fmol of backbone and 68 fmol of gene fragment in 20 μl reactions (New England Biolabs, Ipswitch, MA). The reactions were incubated for 15 min at 50 °C. 3 μl of reaction mix were transformed into *Mix & Go!* Competent Cells - DH5 Alpha (Zymo Research Corporation, Irvine, CA). Correct assembly was checked by colony PCR followed by Sanger sequencing of the gene fragment region and the Gibson junctions. The sequence-confirmed plasmid was named pDL00211. Afterwards, the plasmid was digested with HindIII-HF (New England Biolabs, Ipswitch, MA) and SacI-HF (New England Biolabs, Ipswitch, MA) and treated with Quick CIP alkaline phosphatase (New England Biolabs, Ipswitch, MA). A fragment containing the CYC terminator, originating from plasmid pGD009 ^20^ was ligated into the backbone using NEB T4 Ligase (New England Biolabs, Ipswitch, MA) according to the manufacturers protocol using 29 fmol of backbone and 87 fmol of insert. Correct assembly was checked by colony PCR followed by Sanger sequencing of the CYC terminator and the backbone regions up- and downstream of the restriction sites. The sequence-confirmed plasmid was named pDL00212.

For the prey intermediate plasmid, a gene fragment was ordered from Twist carrying an SP1 Illumina primer binding site, a 24 bp placeholder barcode sequence (AAGTTCGTTGCATCACCTAGCCAA; prey barcode), a 188 bp random spacer sequence, an SP2 Illumina primer binding site, the CYC promoter, a 3 x Flag-tag fused to a DHFR C-terminal half (from here on referred to as FR-tag), a Linker sequence (Linker L4, GASGSAAGGSGSAGSGASAS), the JUN bZIP wt sequence (consisting of the wt DNA-binding domain (DBD) and the five heptads of the wt zipper domain), and Linker L3 was cloned into the pUC19 backbone which had been digested with HindIII-HF (New England Biolabs, Ipswitch, MA) and EcoRI (New England Biolabs, Ipswitch, MA) via Gibson assembly using 62.2 fmol of backbone and 121.4 fmol of gene fragment in 20 μl reactions. The reactions were incubated for 15 min at 50 °C and 3 μl were transformed into *Mix & Go!* Competent Cells - DH5 Alpha (Zymo Research Corporation, Irvine, CA). Correct assembly was checked by colony PCR followed by Sanger sequencing of the gene fragment region and the Gibson junctions. The sequence-confirmed plasmid was named pDL00210.

### Experimental design

The JUN variant library was constructed by overlap-extension PCR as described in Diss and Lehner, 2018. In contrast to the cited library, the JUN sequences were truncated by three amino acids located C-terminally of the fifth heptad. These amino acids are not part of the heptad repeats and were thus not mutagenized in the original design. Here, they were completely removed from the construct. The resulting leucine zipper domain is thus 62 amino acid long, with 27 amino acids for the DNA binding domain and 35 amino acid for the zipper domain, of which only the first 32 where mutated for technical reasons. In the final library, JUN variants were cloned into the prey site of the bindingPCA plasmid (fused to the FR-tag) and the 54 wt bZIP sequences were cloned into the bait site (fused to the DH-tag).

### Barcoding strategy

Every bait-prey pair was identifiable by a double barcode where the bait barcode had a length of 20 bp and the prey barcode had a length of 24 bp. The two barcodes were separated by a 6 bp restriction site so that the full double barcode could be sequenced using the Illumina HiSeq single-end 50 bp sequencing run type. The barcodes were randomly generated using primers with N_20_- and N_24_- overhangs, respectively. Different strategies were used during cloning of the bait and prey intermediate libraries which will be described in detail below.

### Cloning of barcoded bait intermediate plasmids

The 54 bait intermediate plasmids were first cloned individually in pGD009 ^20^ from human cDNA or ORFeome clones and verified by Sanger sequencing. Restriction sites in the CREM and DDIT3 sequences were removed by targeted mutagenesis to a synonymous codon.

To clone them in the bait intermediate plasmid pDL00212 and barcode them, four building blocks needed to be prepared: 1) the backbone by digesting pDL00212 with BstEII-HF (New England Biolabs, Ipswitch, MA) and BamHI-HF (New England Biolabs, Ipswitch, MA) and de-phosphorylating it with Quick CIP alkaline phosphatase (New England Biolabs, Ipswitch, MA); 2) the CYC terminator by digesting pDL00212 with HindIII-HF (New England Biolabs, Ipswitch, MA) and SpeI-HF (New England Biolabs, Ipswitch, MA); 3) the wt bZIP sequences by digesting them out of them original plasmids with BamHI-HF (New England Biolabs, Ipswitch, MA) and SpeI-HF (New England Biolabs, Ipswitch, MA;, and 4) the barcode fragment digested with BstEII-HF (New England Biolabs, Ipswitch, MA) and HindIII-HF (New England Biolabs, Ipswitch, MA). The barcode fragment was created by PCR. The oligonucleotide oDL00549 containing a 20 bp stretch of randomly generated nucleotides (N_20_) was made double stranded using oligonucleotides oDL00122 and oDL00551. A 100 μl setup using 40 pmol of oDL00549 and 200 pmol each of oDL00122 and oDL00551 using the Q5 Master Mix (New England Biolabs, Ipswitch, MA) was run with an initial denaturation at 98 °C for 30 sec, 4 cycles of 10 sec denaturation at 98 °C, 30 sec annealing at 55 °C, and 15 sec elongation at 72 °C, followed by a final extension for 1.5 min at 72 °C. The product was treated with ExoSAP-IT (Thermo Fisher Scientific, Waltham, MA) to remove residual single-stranded oligonucleotides and afterwards purified with the QIAGEN Mini Elute Reaction Cleanup Kit (QIAGEN, Hilden, Germany). The purified product was then digested with BstEII-HF (New England Biolabs, Ipswitch, MA) and HindIII-HF (New England Biolabs, Ipswitch, MA) and afterwards gel-purified using the QIAquick Gel Extraction Kit (QIAGEN, Hilden, Germany) to be ready for cloning. The final plasmids were assembled using NEB T4 Ligase (New England Biolabs, Ipswitch, MA) according to the manufacturers protocol with ca. 14.5 fmol of backbone and 29 fmol of each of the other ingredients. The resulting clones were confirmed via Sanger-sequencing. For each ligation, one clone with a distinctive barcode was chosen (Table S1)

### Cloning of barcoded prey intermediate libraries using mutagenic overlap-extension PCR

The JUN intermediate library was constructed in two steps. The variant library was first cloned into pDL00212, and barcodes were then inserted. To create the variant library, mutagenic overlap-extension PCR was performed as previously described ^20^. However, instead of cloning the library into the backbone via Gibson assembly, the fragments as well as pDL00212 were digested with NheI-HF (New England Biolabs, Ipswitch, MA) and HindIII-HF (New England Biolabs, Ipswitch, MA). The backbone was de-phosphorylated with Quick CIP alkaline phosphatase (New England Biolabs, Ipswitch, MA). All components were ligated together using NEB T4 Ligase (New England Biolabs, Ipswitch, MA) according to the manufacturers protocol. The reactions were performed in a 20 μl reaction using 21.9 fmol of backbone and 43.6 fmol of insert. The reaction was incubated overnight in a PCR cycler using the temperature-cycle ligation (TCL) protocol. This protocol consists of 4 cycles of 10 min at 22 °C and 10 min at 16 °C, followed by 49 cycles of 30 sec each at 4 °C, 27 °C, and 13 °C, then 10 cycles of 30 sec at 4 °C, 1 hour at 16 °C, 10 min at 22 °C, and 10 min at 16 °C, another 49 cycles of 30 sec each at 4 °C, 27 °C, and 13 °C, followed finally by 30 sec at 4 °C, 1 hour at 16 °C, 1 hour at 22 °C and 3 h at 18 °C. Afterwards, the reaction was dialyzed against MilliQ H_2_O (Merck Millipore, Darmstadt, Germany) to reduce the salt concentration using nitrocellulose membranes with 0.025 μm pores (Merck Millipore, Darmstadt, Germany) and then concentrated down to 5 – 10 μl in a speedvac. The concentrated ligated products were then transformed into NEB 10-beta Electrocompetent *E.coli* (New England Biolabs, Ipswitch, MA). Three transformations were performed to increase the yield. For each transformation, 1.5 μl of concentrated ligation reactions was mixed with 25 μl of competent cells and pipetted into a pre-chilled 0.1 cm Gene Pulser Cuvette (Bio-Rad, Hercules, California). Each mixture was electroporated using a Gene Pulser Xcell (BioRad, Hercules, California) using the exponential protocol with 2.0 kV, 200 Ω and 25 μF. After electroporation, the cells were immediately re-suspended in 488 μl of SOC medium that had been pre-warmed to 37 °C. The cuvette was rinsed with another 488 μl of that same medium. All three transformations were pooled and incubated for 30 min at 37 °C. Afterwards, 1 μl of recovered cells was diluted 1/1,000, and 10 μl and 100 μl of this dilution were plated on LB + 2X Amp petri dishes and incubated for 16 h at 37 °C. These plates were used to quantify the number of transformants per transformation. The remaining cells were poured in 150 mL of LB + 4X Amp and incubated at 37 °C and 180 rpm shaking for 16 h. In the morning, the colonies were counted and a total of 23 Mio transformants were obtained. 700ng of the plasmid library was isolated from the liquid culture using the NucleoBond® PC 500 kit (Macherey-Nagel, Düren, Germany).

To barcode the library, a barcode fragment was generated by PCR-amplifying a 326 bp amplicon from the pDL00210 template and using the Q5 High-Fidelity 2X Master Mix (New England Biolabs, Ipswitch, MA) and primers oDL00747 and oDL00748. Primer oDL00747 anneals just downstream of the barcode placeholder sequence and contains a 24 bp overhang of random bases (N_24_) in place of the barcode sequence. The PCR was run for 30 cycles using an annealing temperature of 52 °C for 30 sec and an extension temperature of 72 °C for 1 min. The resulting amplicon was cut from a 1 % agarose gel and purified using the QIAGEN PCR purification kit (QIAGEN, Hilden, Germany). Afterwards, the purified amplicon was digested with AvrII (New England Biolabs, Ipswitch, MA) and XhoI (New England Biolabs, Ipswitch, MA) overnight with an excess of restriction enzymes and gel-purified once more. The resulting fragment was then cloned into the library that had been digested with AvrII (New England Biolabs, Ipswitch, MA) and XhoI (New England Biolabs, Ipswitch, MA) and de-phosphorylated with Quick CIP alkaline phosphatase (New England Biolabs, Ipswitch, MA). The ligation was performed using NEB T4 ligase (New England Biolabs, Ipswitch, MA) according to the manufacturers protocol with 23 fmol of backbone and 46 fmol of barcode insert. The reaction mix was incubated overnight using the TCL protocol as described above, followed by dialysis and concentration also as described above. The transformation into NEB 10-beta electrocompetent *E.coli* (New England Biolabs, Ipswitch, MA) was performed slightly differently. Since the number of barcodes per variant was aimed to be restricted to an average of 30, only 30,000 transformants were supposed to be harvested. A single transformation was performed and then plated for colony counting as described above. The remaining cells were used to inoculate 12 times 10 mL of LB + 4X Amp using 2 x 5 μl, 2 x 10 μl, 2 x 15 μl, 2 x 20 μl, 2 x 50 μl, and 2 x 100 μl of recovery culture. In the morning, the amount of transformants was quantified by counting colonies on the plates and the number of liquid cultures containing a total of ca. 30,000 transformants was pooled and purified by splitting the entire pellet over 10 Miniprep columns (QIAGEN, Hilden, Germany) and eluting each in 30 μl of EB Buffer. This resulted in 145 μg of plasmid library. Successful barcoding was assessed via Sanger sequencing of eight clones. All but one clone showed a correct barcode insertion.

### Barcode-variant association

To put the variant and barcode in close proximity and ensure a sequencing amplicon size suitable for Illumina sequencing, the intermediate library was digested with I-SceI (New England Biolabs, Ipswitch, MA) and NheI-HF (New England Biolabs, Ipswitch, MA) to remove parts of the spacer sequence, the SP2 Illumina primer binding site, the CYC promoter, and the 3x Flag tag – FR fragment – L4 fusion. Afterwards, the ends were blunted with NEB DNA Polymerase I, Large (Klenow) Fragment (New England Biolabs, Ipswitch, MA) according to the manufacturers protocol and then purified using the QIAGEN Mini Elute Reaction Cleanup Kit (QIAGEN, Hilden, Germany) and eluted with 16 μl of buffer EB. The blunt-ended fragments were then re-circularized using NEB T4 ligase (New England Biolabs, Ipswitch, MA). Five reactions of 50 μl with 19 fmol of fragment each were setup and incubated overnight at 15 °C. In the morning, 30 μl per reaction were dialyzed as described above. Afterwards, the five reactions were pooled and concentrated on the speedvac to ca. 20 μl. A single transformation was performed into NEB electrocompetent bacteria (New England Biolabs, Ipswitch, MA) as described above. The non-plated cells were poured in 50 mL of LB + 4x Amp and incubated at 37 °C for 16 h. In the morning, a total of ca. 70 Mio transformants were counted. The cells were harvested, and 230 μg of DNA was extracted using the NucleoBond® PC 500 kit (Macherey-Nagel, Düren, Germany). Efficiency of the re-circularization was checked by Sanger sequencing of eight clones. All showed the expected results, with the barcode and the JUN variant sequence separated by 52 bp.

To prepare the library for sequencing, a two-step PCR was performed to add the Illumina adapter sequences to the amplicons. Since only the SP1 primer binding site was still present in the plasmid, the SP2 primer binding site had to be added via the first round of PCR. For this, 2 ng of the re-cloned library were used as template and a 367 bp barcode-variant fragment was amplified using the KAPA HiFi HotStart DNA Polymerase Kit (Roche Sequencing Solutions Inc, Pleasanton, CA) and primers oDL00345 and oDL00518. The PCR was run for 10 cycles, the annealing was performed at 64 °C for 15 sec and the extension at 72 °C for 15 sec. Afterwards, the reaction was incubated with 20 μl of ExoSAP-IT (Thermo Fisher Scientific, Waltham, MA) at 37 °C for 15 min to remove residual oligonucleotides, followed by heat inactivation at 80 °C for 15 min. The reaction was then purified using the QIAGEN Mini Elute Reaction Cleanup Kit (QIAGEN, Hilden, Germany) and eluted with 12 μl of buffer EB. For the second round of PCR, the entire eluate was used as input for a 300 μl PCR reaction mastermix with the KAPA HiFi HotStart DNA Polymerase kit (Roche Sequencing Solutions Inc, Pleasanton, CA) using primers oDL00345 and oDL00346. The mastermix was split in 6 x 50 μl and the PCR was run for another 10 cycles with an annealing temperature of 72 °C. After PCR, the six reactions were pooled. After verifying the specificity of the amplification on gel, the PCR was gel-purified with the QIAquick Gel Extraction Kit (QIAGEN, Hilden, Germany). The eluate was once more quality-checked with a Bioanalyzer (Agilent Technologies, Santa Clara, CA) which showed chromatograms of single peaks at the correct size. For further quality control, a qPCR was performed using the KAPA Library Quantification Standards with Primer (Roche Sequencing Solutions Inc, Pleasanton, CA). The library was then sequenced on an Illumina NextSeq 500 Sequencing System (Illumina, San Diego, CA) with the Mid-output 300 cycle paired-end sequencing run type. To deal with the low complexity of the library, 30 % of PhiX phage library were spiked in.

Adapters were first removed using cutadapt 1.18^44^. The 5’ adapter was removed from the forward read upstream and including the AvrII site, while the 5’ adapter was removed from the reverse read downstream and including the HindIII and BamHI sites present in tandem. This was done using the default error rate of 0.1. The last base, which seems to be systematically added by the sequencer, was removed from both forward and reverse read. The minimum overlap was set to 6 and max number of N per read set to 0. Forward and reverse reads were then merged using PEAR 0.9.11^45^ with a minimum overlap size of 12, min and max read length of 249 and 259 respecitvely (expected length is 254) and a p-value of 0.05. Merged reads where then pre-processed in R using the shortread package. First, the read was split between barcode and variant sequences. For the barcode, the first 24 bp of the read were used since the sequence upstream of the expected barcode positoon was discarded by cutadapt. Even if there is an indel, these are the 24 bp that will correspond to the barcode during the deepPCA. For the variant, a perfect and unique match of the 10 bp directly upstream of the first JUN codon was required and the sequence downstream is used as the JUN variant sequence since all sequences downstream of the stop codon were discarded by cutadapt.

Read counts were then aggregated to unique barcode-variant sequence pairs. To identify true unique barcode-variant pairs present in the sample from sequencing errors or PCR template switching, which should be at lower frequency, we also counted the total counts for individual barcode and variant sequences. Each unique barcode-variant pair was then also expressed as the percentage of the corresponding total barcode and variant counts. Barcode-variant pairs with less than 10 reads were filtered out since they are far from the 1000 read average expected for the 30,000 transformant obtained and thus likely to be sequencing errors. This reduced the number of unique barcode sequences from ∼500k to 30416, indeed close from our expectation. Next, read counts that likely correspond to sequencing error of the same barcode were collapsed. Thanks to the barcode length, it is highly unlikely the barcodes close in sequence were cloned to the same variant. For each unique variant, read counts from all barcodes with a Hamming distance of 4 bp or less were collapsed to the barcode with the highest frequency. This was then repeated to the second highest frequency barcode, and iteratively until all remaining barcode sequences were processed and all true barcodes associated to this variant were identified. This step further reduced the number of unique barcode sequences to 28969. Plotting read counts of barcode-variant pairs as a function of both total barcode and variant counts reveals a cluster of pairs where both the barcode and the variant have a high total read count but the pair has a low read count. These are likely chimeras that result from template switching during PCR, and were removed by filtering out pairs with less than 100 counts for which the total barcode and variant counts were higher than 300 and 3000 respectively. Finally, for each unique barcode, variant read counts that correspond to sequencing errors were collapsed to the highest frequency variant. A lower frequency variant was considered a sequencing error if it had a Hamming distance of not more than 1 compared to the highest frequency variant, if this mutation was in a different codon and if its total count is 100x lower than the total count of the highest frequency one. If the highest frequency variant was the wild-type JUN sequence, then the variants from which to collapse the count also could not be a variant expected from the mutagenesis design. After this process, unique barcodes for which all associated variants could not be collapsed to the highest frequency one were flagged as promiscuous. 167 barcodes were also flagged as having a Hamming distance <= 4 to at least one other barcode and for which sequencing error during deepPCA could falsely attribute the read between two closely related barcodes. Ultimately, we obtained a total of 28336 high confidence unique barcodes out of which 23385 were associated to expected JUN variant sequences for an average of 38 barcodes per amino acid variant.

### Final library cloning

To prepare the backbone, all wild-type bZIP intermediate plasmids were digested with AvrII (New England Biolabs, Ipswitch, MA) and HindIII-HF (New England Biolabs, Ipswitch, MA), dephosphorylated with Quick CIP alkaline phosphatase (New England Biolabs, Ipswitch, MA), gel-purified with the QIAquick Gel Extraction Kit (QIAGEN, Hilden, Germany) and pooled at equimolar ratios. To prepare the inserts consisting of the Jun variants fused to the FR fragment, the upstream CYC promoter and the variant barcode, the prey intermediate library was digested with AvrII (New England Biolabs, Ipswitch, MA) and HindIII-HF (New England Biolabs, Ipswitch, MA) and gel-purified with the QIAquick Gel Extraction Kit (QIAGEN, Hilden, Germany).

A ligation reaction containing 200 fmol of backbone pool, 600 fmol of insert pool, 10 μl of NEB T4 ligation buffer (New England Biolabs, Ipswitch, MA), 5 μl of NEB T4 ligase (New England Biolabs, Ipswitch, MA) filled up to a volume of 100 μl with MilliQ H_2_O (Merck Millipore, Darmstadt, Germany) was setup. The reaction was split in 2x 50 μl and incubated in a PCR cycler with the TCL program as described above. Afterwards, the ligase was heat-inactivated at 65 °C for 15 min, the 2x 50 μl were pooled and then split in 3 x 33 μl which were each dialyzed against 50 mL of MilliQ H_2_O (Merck Millipore, Darmstadt, Germany) as described above. The dialyzed samples were pooled, which resulted in a total volume of ca. 80 μl. This was concentrated to 25 μl with a speedvac. Two transformations into NEB 10 beta electrocompetent cells (New England Biolabs, Ipswitch, MA) were performed as described above. After recovery, the culture was spread on 35 individual 15 cm diameter LB + 2x Amp plates. The cells were grown to a lawn at 37 °C for approximately 16 h and then scraped off the plates with sterile H_2_O and a plastic spatula. The pellet was washed once with sterile H_2_O and then each library was purified over two columns of the NucleoBond® PC 500 kit (Macherey-Nagel, Düren, Germany). A total of 17 Mio were harvested. The plasmid extractions yielded ca. 1,245 μg of DNA. The quality of the libraries was assessed via test digests, which showed the expected band patterns.

### Large-scale yeast transformation

The transformation was performed in six replicates following a slightly modified protocol from the one described in ^20^. Yeast cells of the BY4742 strain were streaked from a glycerol stock on YPAD plates and incubated at 30 °C for two days prior to the experiment. For each replicate, a single colony was used to inoculate 15 mL of liquid YPAD and grown to saturation overnight at 30 °C, 200 rpm shaking. In the morning, OD600 was measured and for each replicate, a culture of 350 mL pre-warmed YPAD was inoculated at an OD600 of 0.3 and grown at 30 °C and 200 rpm shaking for about 4.5 h until an OD600 of 1.2 to 1.6 was reached. The cells were harvested for 10 min at 3,000 g. Afterwards, the cells were washed with 50 mL H_2_O and centrifuged for 5 min at 3,000 g, the supernatant was discarded, and a second washing step with 50 mL SORB was performed in the same way. Finally, the pellets were re-suspended in 14 mL SORB and incubated for 30 min at RT on a wheel. Next, 350 μl of 10 mg/mL pre-boiled salmon sperm DNA (Agilent Technologies, Santa Clara, CA) were added and mixed thoroughly with the cells before adding 7 μg of the final library to each replicate and mixing by inversion. Each replicate was then split in two times 7 ml and distributed over two 50 mL Falcon tubes. Afterwards, 35 mL of Plate Mixture were added to each tube and the tubes were then incubated for another 30 min at RT on a wheel. Then, 3.5 mL of DMSO (AppliChem, Darmstadt, Germany) were added to each tube, mixed well by inversion, and the tubes were heat-shocked for 20 min in a 42 °C water bath. To homogenize the heat within the tubes, they were inverted 6 times after 1 min, 2 min 30 sec, 5 min, 7 min, 10 min, and 15 min. Afterwards, the tubes were centrifuged for 5 min at 3,000 rpm. The supernatant was removed by pouring first, followed by quick spin and aspiration of the leftover supernatant with a vacuum pump to get rid of any remnants. The pellets were then each re-suspended in 50 mL of pre-warmed recovery medium and incubated at 30 °C for 1 hour without agitation. They were harvested by centrifuging for 5 min at 3,000 g. The two pellets per sample were then pooled again by resuspending in 50 mL SC-ura and harvested once more for 5 min at 3,000 g. After this, the supernatant was discarded, and each pellet was resuspended in 700 mL of SC-ura in 5 L Erlenmeyer flasks. From this culture, 10 μl and 50 μl of each sample were plated on SC-ura plates to assess the transformation efficiency. The plates were incubated at 30 °C for 48 h. The cultures were incubated at 30 °C and 200 rpm shaking for 48 h until they reached saturation. The numbers of transformants varied between 7 and 18 Mio, ensuring at least 10-fold library coverage in each replicate.

### BindingPCA

After the transformant selection culture had reached saturation after about 48 h of growth, a second selection cycle was performed by inoculating a culture of 1 L of SC-ura/ade/met at and OD600 of 0.1 and letting it grow until an OD600 of 1.2 – 1.4. From this culture, a competition culture of 1 L competition medium (SC-ura/ade/met + 200 μg/mL MTX in 2% DMSO) was inoculated at an OD600 of 0.05 and grown for 5 generations until an OD600 of 1.6. The INPUT samples were harvested from the remaining cells of the second selection culture by centrifugation for 5 min at 3,000 g and two washes with 50 mL H_2_O and then frozen at -20 °C. The OUTPUT samples were harvested from the competition cultures in the same way.

### DNA extraction

DNA was extracted from the yeast pellets using a yeast Midiprep protocol to enrich plasmid over genomic DNA. The cells were first spheroblasted by thawing at RT, re-suspending in 20 mL DTT buffer, and incubating at 30 °C and 200 rpm shaking for 15 min. The cells were then harvested at 2,500 g for 5 min, re-suspended in 20 mL Zymolyase buffer and incubated at 30 °C and 200 rpm shaking for 1.5 h. The spheroblasted cells were collected in 50 mL Falcon tubes and spun for 5 min at 2,500 g and re-suspended in 7.5 mL of home-made buffer P1 (according to the manufacturers recipe, QIAGEN, Düren, Germany). 7.5 mL of home-made buffer P2 (according to the manufacturers recipe, QIAGEN, Düren, Germany) were added, the samples were mixed by inverting the tubes several times and incubated at RT for 10 min. 7.5 mL of pre-cooled buffer P3 (QIAGEN, Düren, Germany) were added and the samples were again homogenized by inverting the tubes several times. The samples were centrifuged for 15 min at 3,000 g and 4 °C and the supernatant was then recovered in 50 mL centrifugation tubes (Nalgene, Rochester, NY, USA) and re-centrifuged for 15 min at 15,000 g and 4 °C. The supernatant was then filtered to remove remaining cell debris. The cleared supernatant was applied to columns provided in the Plasmid Midi Kit (QIAGEN, Düren, Germany) that had been equilibrated with 10 mL buffer QBT. Once all of the supernatant had been applied, the columns were washed twice with 10 mL buffer QC and finally, the DNA was eluted with 5 mL buffer QF and collected in 15 mL Falcon tubes. 3.5 mL of Isopropanol was added to each sample and mixed. The samples were then distributed evenly over 2 x 5 mL reaction tubes (Eppendorf, Hamburg, Germany) and spun at 14,200 g for 15 min. The supernatant was removed and the two pellets per sample were pooled by re-suspending each in 500 mL freshly prepared 70 % ethanol and pipetting them together in a normal 1.5 mL reaction tube. The pellets were spun down again at 13,000 rpm for 1 min and then left to dry at RT. Finally, the pellets were re-suspended in 100 μl of buffer EB.

To determine the molar concentration of the plasmid in each sample, as well as the enrichment over genomic DNA, a qPCR was performed using primers OGD241 and OGD242 that bind in the plasmid backbone. For this, the extracts were diluted 1/200 and a standard curve was made using clean plasmid originating from a Miniprep that had been made of one clone of the library and of which the concentration had been determined using the Qubit system (Invitrogen, Waltham, MA, USA). The concentrations used for the standard curve were 0.4 ng/μl, 0.08 ng/μl, 0.016 ng/μl, 0.0032 ng/μl, 0.00064 ng/μl, 0.000128 ng/μl, and 0.0000256 ng/μl. The qPCR was performed using the 2X SsoAdvanced Univeral SYBR Green Supermix (Bio-Rad Laboratories, Hercules, CA, USA) according to the manufacturers protocol.

### Sequencing library preparation

Both Illumina primer binding sites (SP1 and SP2) are already present in the plasmid, so only a single round of PCR had to be performed to prepare the final sequencing libraries. 400 Mio molecules of plasmid for each sample were used as template to ensure at least 10-fold coverage of input molecules over expected read counts. To enable multiplexed sequencing, primers of the NEBNext dual primer set 1 (New England Biolabs, Ipswitch, MA) were used. The INPUT samples used the NEBNext i501 primer as forward primer and the NEBNext i701 to i706 primers as reverse primers. The OUTPUT samples used the NEBNext i502 primer as forward primer and also the NEBNext i701 to i706 primers as reverse primer. The libraries were amplified with Q5 polymerase (New England Biolabs, Ipswitch, MA) in a 50 μl reaction using 400 Mio template molecules, 0.25 μM of each primer, 200 μM dNTPs (New England Biolabs, Ipswitch, MA), 1X concentrated Q5 reaction buffer (New England Biolabs, Ipswitch, MA) and 0.5 μl polymerase. The PCR was run for 18 cycles with 63 °C annealing for 30 sec and 72 °C extension for 30 sec. Successful PCR was confirmed by checking 5 μl of each sample on a 1% Agarose gel. The rest of each reaction was then loaded on a separate 1 % Agarose gel, and the 375 bp amplicon were cut out and purified using the MinElute Gel Extraction kit (QIAGEN, Düren, Germany) and eluted in 20 μl buffer EB. The concentrations were measured using the Qubit system (Invitrogen, Waltham, MA, USA). The concentration of individual samples was confirmed using the KAPA Library Quantification Standards with Primer (Roche Sequencing Solutions Inc, Pleasanton, CA). All 6 INPUT and 6 OUTPUT samples per screen were pooled at equimolar ratios. The concentration of the final pool was confirmed once more first with Qubit, then with qPCR. A final quality control was performed by loading the library on a Bioanalyzer (Agilent Technologies, Santa Clara, CA). Finally, the library was submitted for sequencing on a single lane of the Illumina HiSeq 2500 System (Illumina, San Diego, CA) with the 50 bp single-end sequencing run type. To increase sequencing quality and clustering efficiency, 25 % of PhiX phage library were spiked into the library.

### Sequencing data processing and binding score calculation

Raw sequencing reads were filtered for an average Q score >= 20, split into the wild-type barcode (first 20bp) and the variant barcode (last 24bp), and collapsed to obtain read counts for each barcode pair. Wildtype barcode reads with a Hamming distance <= 2 to the expected true barcode sequence were kept. Jun variant barcode reads with a Hamming distance < 2 to the expected true barcodes and a difference in Hamming > 2 between the two best matches were kept. Read counts of barcode pairs matching the same combination of JUN amino acid substitution and wild-type partners were collapsed. Output samples were sequenced twice to increase coverage, and the read counts of the sample were added. Read count tables were then processed with DiMSum v1.3^23^ (https://github.com/lehner-lab/DiMSum) using STEAM stages 4-5 (“countPath” option) and setting the JUN wild-type x FOS pair as an arbitrary reference to obtain binding scores and associated errors.

### Thermodynamic model

We used MoCHI (https://github.com/lehner-lab/MoCHI)^21,22^ to fit a global mechanistic model to the bPCA data where JUN variants binding to all partner bZIPs are modelled as an equilibrium between two states: unfolded and unbound or folded and bound.

We assume that free energy changes of binding are additive and shared between all partner bZIPs i.e. the total binding free energy changes of an arbitrary variant to an arbitrary bZIP relative to the wild-type JUN sequence is simply the sum over residue-specific energies corresponding to all constituent single amino acid substitutions.

In order to model bZIP-specific modifications in binding affinity we dummy encoded the presence/absence of each partner bZIP in a 51-mer amino acid sequence appended to the JUN variant amino acid sequence where a A>C substitution denotes the presence of the corresponding partner bZIP. The associated inferred free energy changes for these dummy substitutions correspond to additive bZIP-specific binding affinity effects.

However, for some partners this thermodynamic model did not provide a good fit to the bPCA data and suggested a partner-dependent non-linear trend corresponding to a multiplicative energy term (effectively controlling the steepness of the sigmoid gradient). We therefore modified the model to infer one such term for each bZIP partner in addition to the additive effects described above. This modification indeed reduced the bias observed in the residuals, although without significantly improving the overall goodness of the fit (R^2^ = 0.91 in both cases). Interestingly, these multiplicative terms are equivalent to Hill’s coefficient, which is determined by the cooperativity of the interaction between multiple subunits, although the underlying mechanisms in the context of this assay remains speculative pending further investigation.

We configured MoCHI parameters to specify a neural network architecture consisting of two additive trait (free energy) layers i.e. one corresponding to the additive JUN and dummy-encoded bZIP partner binding free energy changes (“Binding”) and another for the bZIP-specific multiplicative effects (“BindingMod”). A single linear transformation layer outputs the predicted bPCA binding score. The specified non-linear transformation “TwoStateFractionFoldedMod” derives from the Boltzmann distribution function relate binding energies to proportions of folded and bound molecules.

A random 30% of variant-partner combinations was held out during model training, with 20% representing the validation data and 10% representing the test data. Validation data was used to evaluate training progress and optimise hyperparameters (batch size). Optimal hyperparameters were defined as those resulting in the smallest validation loss after 100 training epochs. Test data was used to assess final model performance.

Models were trained with default settings i.e. for a maximum of 1000 epochs using the Adam optimization algorithm with an initial learning rate of 0.05. MoCHI reduces the learning rate exponentially (γ = 0.98) if the validation loss has not improved in the most recent ten epochs compared to the preceding ten epochs. In addition, MoCHI stops model training early if the wild-type free energy terms over the most recent ten epochs have stabilised (standard deviation 10^-3^).

### Comparison to in vitro measurements

In vitro FRET measurements were obtained from Reinke et al^24^. K_D_ measured at 37C were converted to DG, normalized to the JUN-FOS DG to obtain DDG and the Pearson correlation with the partner’s DDG fitted by MoCHI was calculated.

### Amino acid features predictive of global energy

To determine which amino acid feature relates to global energy, we fitted a linear model between a database of >500 amino acid features^26^ and the DDG fitted by MoCHI. We retained only the 536 features for which values for all 20 amino acids were available. We performed a forward feature search by an iterative process where each feature was used one-by-one together with the heptad number and the position within the heptad of each mutant. The best predictive feature was retained, and the search was repeated including the previously identified predictive feature. We next determined the best feature at each heptad position. The data was split by heptad position, and at each position, each feature was tested alone and the best one was retained.

### Decomposing mutational effect into global and specific effects

Despite the multiplicative term, the fit for ATF4 was biased and the fitted sigmoid does not fit the apparent trend. To not bias the analysis of specific effects, all measurements of binding to ATF4 were therefore filtered out. Additionally, all measurements with 10 or less counts in any input replicate or 0 count in any output replicates were filtered out because of the high error inherent to low read counts. Mutational effect was then calculated by subtracting the binding score of interaction between wild-type JUN and each corresponding partner. The measurement error of the mutational effect was calculated by propagating the error of the mutant and the corresponding wild-type pair, and a p-value was calculated by performing a one-sample t-test. P-values were adjusted by the FDR method.

Mutational effects due to global energy effects were calculated by subtracting the binding score predicted by MoCHI from the binding score of the corresponding wild-type JUN – partner combination. Corresponding errors were calculated by propagating the measurement and prediction errors, p-values were calculated by a one-sample t-test and corrected by the FDR method. Specific effects were calculated by subtracting the binding score predicted by MoCHI from the one measured, and error, p-values and FDR calculated as above.

### AlpaFold2 predictions

Predictions of protein complexes between JUN and a few partners presented in Fig. 7 were generated using Alphafold2-multimer (v2.3.2) using the sequences in Table S1, with multimer models v3 and subsequent model relaxation^46,47^. The pipeline was run through GUIFold^48^ (v0.4). For the generation of multiple sequence alignments, the “full_dbs” protocol was employed. In case of each prediction target, the best model was selected by the smallest inter-subunit average predicted aligned error (PAE). The range of inter-subunit average PAE for all predictions was between 5.5 and 10.0 Å.

## Data availability

Deep sequencing data is available at GEO with accession number GSE245326.

## Code availability

All custom scripts will be provided upon requests

## Acknowledgements

This work was supported by the Novartis Research Foundation (all authors except AJF and BL) and SNF Project grant 197593 (GD, AMB). Work in the lab of BL is funded by the European Research Council (ERC) Advanced grant (883742), the Spanish Ministry of Science and Innovation (LCF/PR/HR21/52410004, EMBL Partnership, Severo Ochoa Centre of Excellence), the Bettencourt Schueller Foundation, the AXA Research Fund, Agencia de Gestio d’Ajuts Universitaris i de Recerca (AGAUR, 2017 SGR 1322), and the CERCA Program/Generalitat de Catalunya. AJF. was funded by a Ramón y Cajal fellowship (RYC2021-033375-I) financed by the Spanish Ministry of Science and Innovation (MCIN/AEI/10.13039/501100011033) and the European Union (NextGenerationEU/PRTR).

## Author contributions

GD and BL conceptualized the project. GD designed the experiments. AMB performed all the experimental work with help from DK and KS. GD analysed the data. AJF fitted the thermodynamic model. GK and SC performed the AlphaFold2 structure predictions. GD and BL interpreted the results and wrote the manuscript.

## Figures

**Fig. S1: Specificity plots for all 52 partners**. Specifcity plots representing the binding scores as a function of the global binding energy (ΔΔG) inferred by MoCHI. The sigmoid curve represents binding scores predicted by MoCHI, and the vertical residual from the curve thus represent the specific effect of this mutation on the interaction with this partner. The three characters in the mutant name corresponds to heptad, position within the heptad and amino acid substitution, respectively. Mutant names are colored according to their heptad position, which further highlights that several mutations at the same few positions have specific effects.

## Notes

### Competing Interest Statement

The authors have declared no competing interest.

## References

1. Crick, F. H. C. The packing of α-helices: simple coiled-coils. Acta Crystallogr. 6, 689– 697 (1953).

2. O’Neil, K. T. & DeGrado, W. F. A thermodynamic scale for the helix-forming tendencies of the commonly occurring amino acids. Science (80-.). 250, 646–651 (1990).

3. O’Shea, E. K., Rutkowski, R. & Kim, P. S. Evidence that the leucine zipper is a coiled coil. Science (80-.). 243, 538–542 (1989).

4. Landschulz, W. H., Johnson, P. F. & McKnight, S. L. The Leucine Zipper: A Hypothetical Structure Common to a New Class of DNA Binding Proteins. Science (80-.). 240, 1759–1764 (1988).

5. Thompson, K. S., Freire, E. & Vinson, C. R. Thermodynamic Characterization of the Structural Stability of the Coiled-Coil Region of the bZIP Transcription Factor GCN4. Biochemistry 32, 5491–5496 (1993).

6. Vinson, C. R., Hai, T. & Boyd, S. M. Dimerization specificity of the leucine zipper-containing bZIP motif on DNA binding: Prediction and rational design. Genes Dev. 7, 1047–1058 (1993).

7. Arndt, K. M., Pelletier, J. N., Müller, K. M., Plückthun, A. & Alber, T. Comparison of in vivo selection and rational design of heterodimeric coiled coils. Structure 10, 1235– 1248 (2002).

8. Mason, J. M., Schmitz, M. A., Müller, K. M. & Arndt, K. M. Semirational design of Jun-Fos coiled coils with increased affinity: Universal implications for leucine zipper prediction and design. PNAS 103, 8989–8994 (2006).

9. Mason, J. M., Müller, K. M. & Arndt, K. M. Positive Aspects of Negative Design: Simultaneous Selection of Specificity and Interaction Stability †. Biochemistry 46, 4804–4814 (2007).

10. Grigoryan, G. & Keating, A. E. Structure-based prediction of bZIP partnering specificity. J. Mol. Biol. 355, 1125–1142 (2006).

11. Grigoryan, G., Reinke, A. W. & Keating, A. E. Design of protein-interaction specificity affords selective bZIP-binding peptides. Nature 458, 859–864 (2009).

12. Reinke, A. W., Grant, R. A. & Keating, A. E. A synthetic coiled-coil interactome provides heterospecific modules for molecular engineering. J. Am. Chem. Soc. 132, 6025–6031 (2010).

13. Potapov, V., Kaplan, J. B. & Keating, A. E. Data-Driven Prediction and Design of bZIP Coiled-Coil Interactions. PLoS Comput. Biol. 11, 1–28 (2015).

14. Ljubetič, A. et al. Design of coiled-coil protein-origami cages that self-assemble in vitro and in vivo. Nat. Biotechnol. 35, 1094–1101 (2017).

15. Lebar, T., Lainšček, D., Merljak, E., Aupič, J. & Jerala, R. A tunable orthogonal coiled-coil interaction toolbox for engineering mammalian cells. Nat. Chem. Biol. 16, 513–519 (2020).

16. Newman, J. R. S. & Keating, A. E. Comprehensive identification of human bZIP interactions with coiled-coil arrays. Science (80-.). 300, 2097–2101 (2003).

17. Pelletier, J. N., Campbell-Valois, F.-X. & Michnick, S. W. Oligomerization domain-directed reassembly of active dihydrofolate reductase from rationally designed fragments. PNAS 95, 12141–12146 (1998).

18. Freschi, L., Torres-Quiroz, F., Dubé, A. K. & Landry, C. R. qPCA: a scalable assay to measure the perturbation of protein-protein interactions in living cells. Mol. Biosyst. 9, 36–43 (2013).

19. Levy, E. D., Kowarzyk, J. & Michnick, S. W. High-resolution mapping of protein concentration reveals principles of proteome architecture and adaptation. Cell Rep. 7, 1333–1340 (2014).

20. Diss, G. & Lehner, B. The genetic landscape of a physical interaction. Elife 7, 1–31 (2018).

21. Faure, A. J. et al. Mapping the energetic and allosteric landscapes of protein binding domains. Nature 604, 175–183 (2022).

22. Weng, C., Faure, A. J. & Lehner, B. The energetic and allosteric landscape for KRAS inhibition. bioRxiv 2022.12.06.519122 (2022). doi:10.1101/2022.12.06.519122

23. Faure, A. J., Schmiedel, J. M., Baeza-Centurion, P. & Lehner, B. DiMSum: An error model and pipeline for analyzing deep mutational scanning data and diagnosing common experimental pathologies. Genome Biol. 21, 1–23 (2020).

24. Reinke, A. W., Baek, J., Ashenberg, O. & Keating. Networks of bZIP Protein-Protein Interactions Diversified Over a Billion Years of Evolution. Science (80-.). 340, 730–735 (2013).

25. Chen, S. et al. The emerging role of XBP1 in cancer. Biomed. Pharmacother. 127, 110069 (2020).

26. Kawashima, S. et al. AAindex: amino acid index database, progress report 2008. Nucleic Acids Res. 36, D202–5 (2008).

27. Qian, N. & Sejnowski, T. J. Predicting the secondary structure of globular proteins using neural network models. J. Mol. Biol. 202, 865–884 (1988).

28. Roseman, M. A. Hydrophobicity of the peptide C=O…H-N hydrogen-bonded group. J. Mol. Biol. 201, 621–623 (1988).

29. Hutchens, J. O. Handbook of Biochemistry. (1970).

30. DeGrado, W. F. & Lear, J. D. Induction of peptide conformation at apolar water interfaces. 1. A study with model peptides of defined hydrophobic periodicity. J. Am. Chem. Soc. 107, 7684–7689 (1985).

31. O’Shea, E. K., Rutkowski, R. & Kim, P. S. Mechanism of specificity in the Fos-Jun oncoprotein heterodimer. Cell 68, 699–708 (1992).

32. Bastolla, U., Porto, M., Roman, H. E. & Vendruscolo, M. Principal eigenvector of contact matrices and hydrophobicity profiles in proteins. Proteins 58, 22–30 (2005).

33. Bigelow, C. C. On the average hydrophobicity of proteins and the relation between it and protein structure. J. Theor. Biol. 16, 187–211 (1967).

34. Nakashima, H. & Nishikawa, K. The amino acid composition is different between the cytoplasmic and extracellular sides in membrane proteins. FEBS Lett. 303, 141–146 (1992).

35. Meirovitch, H., Rackovsky, S. & Scheraga, H. A. Empirical Studies of Hydrophobicity. 1. Effect of Protein Size on the Hydrophobic Behavior of Amino Acids. Macromolecules 13, 1398–1405 (1980).

36. Ponnuswamy, P. K., Prabhakaran, M. & Manavalan, P. Hydrophobic packing and spatial arrangement of amino acid residues in globular proteins. Biochim. Biophys. Acta 623, 301–316 (1980).

37. Podgornaia, A. I. & Laub, M. T. Determinants of specificity in two-component signal transduction. Curr. Opin. Microbiol. 16, 156–162 (2013).

38. Schreiber, G. & Keating, A. E. Protein Binding Specificity versus Promiscuity. Curr. Opin. Struct. Biol. 21, 50–61 (2011).

39. Lite, T. L. V. et al. Uncovering the basis of protein-protein interaction specificity with a combinatorially complete library. Elife 9, 1–57 (2020).

40. McClune, C. J., Alvarez-Buylla, A., Voigt, C. A. & Laub, M. T. Engineering orthogonal signalling pathways reveals the sparse occupancy of sequence space. Nature 574, 702–706 (2019).

41. Aakre, C. D. et al. Evolving New Protein-Protein Interaction Specificity through Promiscuous Intermediates. Cell 163, 594–606 (2015).

42. Ghosea, D. A., Przydziala, K. E., Mahoneya, E. M., Keating, A. E. & Laub, M. T. Marginal specificity in protein interactions constrains evolution of a paralogous family. PNAS 120, 1–10 (2023).

43. Sikorski, R. S. & Hieter, P. A system of shuttle vectors and yeast host strains designed for efficient manipulation of DNA in Saccharomyces cerevisiae. Genetics 122, 19–27 (1989).

44. Martin, M. CUTADAPT removes adapter sequences from high-throughput sequencing reads. EMBnet.journal 17, (2011).

45. Zhang, J., Kobert, K., Flouri, T. & Stamatakis, A. PEAR: a fast and accurate Illumina Paired-End reAd mergeR. Bioinformatics 30, 614–620 (2014).

46. Jumper, J. et al. Highly accurate protein structure prediction with AlphaFold. Nature 596, 583–589 (2021).

47. Evans, R. et al. Protein complex prediction with AlphaFold-Multimer. bioRxiv 03.101, (2022).

48. Kempf, G. & Cavadini, S. GUIFold - A graphical user interface for local AlphaFold2. bioRxiv (2023). doi:10.1101/2023.01.19.521406

49. Glover, J. N. & Harrison, S. C. Crystal structure of the heterodimeric bZIP transcription factor c-Fos-c-Jun bound to DNA. Nature 373, 257–261 (1995).

